# Investigating the causal and consequential role of the nuclear encoded mitochondrial genome in Amyotrophic Lateral Sclerosis

**DOI:** 10.64898/2026.11.10.764216

**Authors:** Aminah T. Ali, Ahmad Al Khleifat, Ammar Al-Chalabi, Alan Hodgkinson, Alfredo Iacoangeli

## Abstract

Mitochondrial dysfunction is implicated in amyotrophic lateral sclerosis (ALS), but which mitochondrial pathways are involved, and whether their involvement is a cause or a consequence of disease remains unclear. We investigated whether nuclear-encoded mitochondrial gene (NEMG) pathways are enriched for ALS genetic risk, which are dysregulated in ALS tissue, whether pathway activity defines patient subgroups that differ in survival, and whether co-expression between the nuclear and mitochondrial genomes is altered.

We tested MitoCarta3.0 pathways for enrichment of ALS genetic risk using MAGMA and the largest European ALS GWAS to date (27,205 cases and 110,881 controls), using hierarchical testing across pathway tiers, and annotated the direction of gene-level effects using published TWAS and SMR results. In >1,500 bulk RNA-seq samples from post-mortem motor cortex and spinal cord (KCL Brain Bank, NYGC ALS Consortium) and iPSC-derived motor neurones (Answer ALS), we performed cell type adjusted gene set enrichment analysis (GSEA), gene set variation analysis (GSVA)-based clustering with exploratory survival analysis, and nuclear-mitochondrial co-expression analysis.

Of seven broad mitochondrial categories, only the mitochondrial central dogma was enriched for ALS risk after correction (P_adj_ = 0.044). Hierarchical testing within this pathway localised the signal to translation (P_adj_ = 0.008) and the mitoribosome (P_adj_ = 0.014). Genetically predicted expression pointed in opposite directions for different mitoribosomal genes (lower *MRPS33* and higher *MRPL28* expression with increased risk), and did not agree in direction with differential expression in ALS tissue more often than expected by chance. A nominal enrichment in ROS and glutathione metabolism was largely attributable to *SOD1*. In tissue, mitochondrial genes were broadly more highly expressed in ALS motor cortex in both cohorts, largely as a global shift across mitochondrial genes. OXPHOS subunits and the mitoribosome were enriched beyond this shift in ALS Consortium motor cortex (both P_FDR_ = 0.027), as were OXPHOS subunits in iPSC-derived motor neurones (P_FDR_ = 2.4 x 10⁻⁸), whereas spinal cord showed no genome-wide enrichment. GSVA based clusters were associated with diagnosis in motor cortex, and nominally with survival in lumbar spinal cord (adjusted HR 0.69, P_unadj_ = 0.013), where cluster structure was weakest. No individual pathway was associated with survival after correction in adjusted models. In all four post-mortem tissues, nuclear-encoded mitochondrial genes were more negatively co-expressed with mtDNA-encoded genes in ALS than in controls, most strongly in lumbar spinal cord, whereas iPSC-derived motor neurones showed the opposite pattern.

In conclusion, inherited ALS risk shows a modest enrichment centred on mitochondrial translation, with gene specific rather than uniform directions of effect. The same pathway, alongside OXPHOS, is more highly expressed in end-stage ALS motor cortex, which may reflect a response to disease, residual differences in cell composition, or both. At the gene level, genetically predicted and observed expression changes did not agree in direction, so whether inherited risk and these tissue changes are linked remains unresolved.

## Introduction

Amyotrophic Lateral Sclerosis (ALS) is a progressive and fatal neurodegenerative disease affecting the upper and lower motor neurones, leading to progressive weakness of voluntary muscles, and eventually death within a few years of symptom onset (Goutman *et al*., 2022b). ALS is thought to be the result of a complex combination of genetics, lifestyle, and environmental exposure (Wang *et al*., 2017), and we are currently able to identify a known genetic cause in approximately 20% of people with ALS (pwALS) (Mehta *et al*., 2022; Iacoangeli *et al*., 2025; Hop *et al*., 2026).

Approximately 10% of individuals with ALS have a family history of the disease, typically attributable to high penetrance mutations in causal genes (Goutman *et al*., 2022a). The remaining 90% are conventionally classified as having “sporadic” ALS, although these individuals may also carry pathogenic variants as a result of the incomplete penetrance of most ALS mutations (Shatunov and Al-Chalabi, 2021; Spargo *et al*., 2022, 2023). In addition, twin studies estimate that the heritability of ALS to be ∼61% (Al-Chalabi *et al*., 2010), suggesting a substantial genetic contribution, even in the absence of a family history. SNP based heritability estimates provide solid evidence of a polygenic component behind disease risk that can play a role in most pwALS (Van Rheenen *et al*., 2021; Hu *et al*., 2026).

The 40+ identified ALS causal genes converge on dysregulating major pathological pathways, including impaired RNA metabolism, altered proteostasis and autophagy, cytoskeletal and axonal transport, DNA repair, and mitochondrial dysfunction (Hardiman *et al*., 2017; Goutman *et al*., 2022a; Nijs and Van Damme, 2024).

Mitochondrial dysfunction is a particularly interesting pathogenic mechanism, with many ALS linked genes, including *CSorf72*, *SOD1*, *TARDBP* and *FUS,* affecting mitochondrial functions either directly or indirectly (Smith, Shaw and De Vos, 2019; Goutman *et al*., 2022a). The intronic GGGGCC hexanucleotide repeat expansion within *CSorf72* (DeJesus-Hernandez *et al*., 2011; Renton *et al*., 2011) is proposed to lead to neurodegeneration through reduced *CSorf72* expression, accumulation of toxic RNA foci, and aggregation of dipeptide repeat proteins (DPRs) (Smeyers, Banchi and Latouche, 2021). Both loss of function and DPR toxicity converge on oxidative phosphorylation. C9orf72 stabilises the complex I assembly factor TIMMDC1 (Wang *et al*., 2021), and poly(GR) binds the complex V subunit ATP5A1 and promotes its degradation (Choi *et al*., 2019). Misfolded mutant SOD1 deposits on the outer membrane of spinal cord mitochondria (Vande Velde *et al*., 2008) and directly inhibits VDAC1, a voltage dependant anion channel (Israelson *et al*., 2010). Mutant TARDBP demonstrates accelerated and increased aggregation (Johnson *et al*., 2009), mislocalisation from the nucleus to the cytoplasm (Barmada *et al*., 2010), and altered RNA splicing (Arnold *et al*., 2013; Fratta *et al*., 2018). TARDBP also enters mitochondria, where it reduces expression of the complex I subunits ND3 and ND6 and causes complex I disassembly (Wang *et al*., 2016). Finally, mutant FUS, also linked to cytoplasmic mislocalisation, aggregation and altered RNA splicing (Kwiatkowski *et al*., 2009; Vance *et al*., 2009; Humphrey *et al*., 2020) binds the ATP synthase subunit ATP5B and disrupts complex V assembly (Deng *et al*., 2018).

The mitochondrial genome itself has also been linked to ALS. Mitochondrial DNA copy number (mtDNA-CN) measured in blood is observed to be higher in pwALS than in controls (Stoccoro *et al*., 2018; Harvey *et al*., 2024), and higher mtDNA-CN was associated with shorter survival (Harvey *et al*., 2024). As mtDNA-CN number is observed to decline with age (Hägg *et al*., 2021; Longchamps *et al*., 2022), Harvey *et al*. suggest that the higher level observed in pwALS is unexpected and represents a response to disease. A possible explanation for this observation comes from work in Parkinson’s disease (PD). In isolated neurones from healthy brain tissue, mtDNA-CN was observed to positively correlate with the number of age related mtDNA deletions, however, this correlation was weaker in neurons from PD brains (Dölle *et al*., 2016).

Dölle *et al*. suggest that this reflects a neuroprotective mechanism allowing neurons to maintain a healthy population of mtDNA, and that this compensatory mechanism is reduced in PD. The raised mtDNA-CN number seen in ALS could represent a similar compensatory response to accumulating mtDNA damage.

Further to this, Stoccoro *et al*., (2018) have reported reduced mitochondrial displacement loop (D-loop) methylation in blood from both presymptomatic and affected SOD1 mutation carriers. The mitochondrial D-loop is a non-coding control region within the mtDNA and is involved in regulating mtDNA replication and transcription. In this study, D-loop methylation was inversely correlated with mtDNA copy number, suggesting a compensatory upregulation of mtDNA (Stoccoro *et al*., 2018). Heteroplasmic mtDNA variation within the D-loop has also been observed to be enriched in post-mortem ALS spinal cord, with in silico analysis suggesting they may alter D-loop secondary structure, and therefore influence replication initiation and transcription (Cox *et al*., 2024). Mitochondrial function is regulated by coordinated expression between the nuclear and the mitochondrial genome (Papier *et al*., 2022), and disruption of this coordination may play a role in the development and progression of ALS.

As suggested by the multiple pathways involved, ALS is heterogenous. Clinically for example, patients differ in site of symptom onset, age at onset and rate of disease progression (Al-Chalabi *et al*., 2016). Heterogeneity is also evident at the molecular level. Studies that integrate patient transcriptomic data from post-mortem motor cortex and spinal cord with machine learning approaches have identified distinct patient subgroups (Tam *et al*., 2019; Eshima *et al*., 2023; Marriott *et al*., 2023; O’Neill *et al*., 2025). Notably, a subgroup relating to oxidative stress and mitochondrial dysfunction is consistently identified across studies and tissues, representing between 25-75% of cases within their respective cohorts (Jammal *et al*., 2026). Because subgroups were derived using different methods, tissues and gene sets, the extent to which they reflect the same underlying mitochondrial defects remains unclear.

A related question is whether mitochondrial dysfunction is a cause, consequence or combination of the two when it comes to motor neuron degeneration. A recent study has shown that FUS- and TARDBP-motor neurons show an early, cell-type-specific disruption of mitochondrial gene expression and axonal mitochondrial transport (Schweingruber *et al*., 2025). In another study of rare, loss of function variants in nuclear-encoded mitochondrial genes, *DNA2* and *ACADM* were associated with ALS survival but not disease risk (Harvey *et al*., 2024).

Defining which mitochondrial pathways are altered, in which patients, and in which order may therefore help identify those most likely to benefit from mitochondria-targeted therapies. This study takes a hypothesis driven approach, restricting analyses to nuclear-encoded mitochondrial genes defined by MitoCarta3.0, to address each of these areas. To identify which pathways are involved upstream, we tested for enrichment of ALS genetic risk. To identify later stage transcriptional changes, we examined post-mortem motor cortex, spinal cord and in patient origin iPSC derived motor neurones. Finally, to ask in which patients, we clustered individuals by mitochondrial pathway activity and tested whether these groups differ in survival outcomes.

## Materials and Methods

### Mitochondrial gene set definitions

Nuclear-encoded mitochondrial genes (NEMGs) and their pathway annotations were defined according to the MitoCarta3.0 database (Rath *et al*., 2021), a curated inventory of 1,136 human protein-coding genes with strong evidence of mitochondrial localisation. Within MitoCarta3.0, genes are hierarchically organised into seven broad functional categories (tier 1), 39 sub-pathways (tier 2) and 103 more specific sets (tier 3), giving 149 pathways in total. These pathway annotations were used to define gene sets in downstream analyses.

### GWAS summary statistics

European-ancestry genome-wide association study (GWAS) summary statistics for ALS, as reported by (Van Rheenen *et al*., 2021) were downloaded from the GWAS catalogue (Cerezo *et al*., 2025) with accession code GCST90027164. These summary statistics come from the largest ALS GWAS conducted to date and include data from 27,205 individuals with ALS and 110,881 individuals without ALS.

### MAGMA gene-set analysis

Gene-set analysis was performed using MAGMA v1.10 (De Leeuw *et al*., 2015) to test whether genetic variants surrounding NEMGs are jointly more strongly associated with ALS than expected by chance. Gene-level association statistics were derived from the (Van Rheenen *et al*., 2021) summary statistics using default MAGMA parameters, with linkage disequilibrium (LD) estimated from the 1000 Genomes Project Phase 3 European reference panel (The 1000 Genomes Project Consortium *et al*., 2015). Gene-level statistics were then regressed on MitoCarta3.0 gene-set membership, with adjustment for gene size, gene density, and minor allele count. One-sided tests were used to assess positive enrichment of ALS association signals within each gene set. Gene-set analyses were performed hierarchically across MitoCarta3.0 categories. First, enrichment was tested across the seven tier 1 functional categories, with correction for seven tests. To identify the biological processes driving enrichment, we tested sub-pathways at tiers 2 and 3, correcting for the number of sub pathways within each parent. For example, the mitochondrial central dogma pathway has three sub-pathways, so was corrected for 3 tests. The mitochondrial central dogma > translation pathway has 5 sub-pathways, so was corrected for 5 tests. Gene sets were mapped to MAGMA gene-level statistics using Entrez gene IDs. Sets with fewer than two mapped genes were not tested, leaving 7, 38 and 96 total sets at tiers 1-3 respectively.

### TWAS and SMR summary statistics

Results from transcriptome-wide association studies (TWAS) and summary-based Mendelian Randomisation (SMR), carried out across multiple independent blood and brain datasets were made available from (Pain *et al*., 2024). TWAS and SMR were performed by integrating the European-ancestry ALS GWAS summary statistics from (Van Rheenen *et al*., 2021), with a comprehensive set of expression quantitative trait loci (eQTL) reference panels spanning multiple brain regions and blood tissues. Brain expression panels included data from GTEx across 13 brain regions, the CommonMind Consortium (CMC) dorsolateral prefrontal cortex (DLPFC), PsychENCODE DLPFC, MetaBrain across five brain regions, and a motor cortex panel derived from the King’s College London (KCL) Brain Bank. Blood expression panels included GTEx, NTR, YFS, and eQTLGen. To confirm that identified associations reflected a shared causal variant rather than linkage, colocalisation analyses were performed using coloc (PP4/(PP3+PP4) > 0.8) for FUSION results and the HEIDI test (p > 0.05) for SMR results. Because PP4/(PP3+PP4) conditions on both traits being associated, we also report the absolute PP4, and regard colocalisation as well supported only where PP4 > 0.5. These analyses use the same GWAS summary statistics as the MAGMA analysis and were used to annotate direction of effect, not as independent replication.

### RNA-seq data preprocessing

Bulk RNA-seq data from three ALS cohorts were included in our study. From the King’s College London and MRC London Neurodegenerative Diseases Brain Bank (KCL Brain Bank; Iacoangeli *et al*., 2021; Jones *et al*., 2021), we obtained 171 post-mortem motor cortex samples (112 ALS, 59 controls), processed as described in (Marriott *et al*., 2023).

From the New York Genome Center ALS Consortium (ALS Consortium; Humphrey *et al*., 2026), we obtained gene-level counts from post-mortem motor cortex and spinal cord. Only samples from donors with ALS or from controls were retained. Donors with ALS-FTD, FTD or other motor neurone diseases were excluded. Medial, lateral and unspecified motor cortex samples were combined into a merged motor cortex set (432 samples from 309 donors). Because some donors contributed samples from more than one motor cortex region, one sample per donor was selected at random, giving 309 samples (270 ALS, 39 controls). Cervical (331 samples; 277 ALS, 54 controls) and lumbar (314 samples; 261 ALS, 53 controls) spinal cord data were analysed separately.

From Answer ALS (Baxi *et al*., 2022), we obtained count data from iPSC-derived motor neurones. Participants classified as having a non-ALS motor neurone disease, or as asymptomatic ALS gene carriers, were excluded, leaving 632 samples with both expression data and metadata (458 ALS, 174 controls).

Covariates were chosen for each dataset according to availability. For the KCL Brain Bank, these were RNA integrity number (RIN), age at death, post-mortem interval, sex, and estimated proportions of astrocytes, neurones, microglia and oligodendrocytes from (Marriott *et al*., 2023). For the ALS Consortium, they were RIN, age at death, sex, site of sample collection, and cell-type proportions estimated by (Humphrey *et al*., 2026). The most abundant cell type in each tissue was omitted, to avoid problems with colinearity. For Answer ALS, covariates were sex, batch, and the percentage of cells staining positive for SMI32, ISL1, NKX6.1, TUJ1, S100β and Nestin. Covariates with a single value within a tissue were excluded from that tissue’s model. Samples with missing values for any covariate were removed. This gave final sample sizes of 171 for the KCL Brain Bank (112 ALS, 59 controls); 269 for ALS Consortium merged motor cortex (237 ALS, 32 controls), 300 for cervical spinal cord (250 ALS, 50 controls) and 288 for lumbar spinal cord (238 ALS, 50 controls); and 542 for Answer ALS (390 ALS, 152 controls).

To capture unmeasured sources of variation, surrogate variables (SVs) were estimated with svaseq from the sva package (Leek, 2014).Counts were normalised for library size using DESeq2 size factors (Love, Huber and Anders, 2014)and restricted to genes with a normalised count of at least 10 in at least 10 samples and a mean normalised count above 1. The full model contained all covariates listed above, including disease status, and the null model contained only an intercept. The number of SVs was estimated with num.sv and capped at (n/4) − p, where n is the number of samples and p the number of parameters in the full model, to keep estimation stable. This yielded 1, 2, 1, 1 and 3 SVs for the KCL Brain Bank, ALS Consortium merged motor cortex, cervical spinal cord, lumbar spinal cord and Answer ALS, respectively.

### Differential gene expression analysis

Differential gene expression (DGE) between ALS and control samples was performed separately for each dataset and tissue. Genes were retained if they had at least 10 reads in at least as many samples as the smaller diagnostic group. Control was set as the reference level, and continuous covariates were included as fixed effects.

For the KCL Brain Bank, Answer ALS, and ALS Consortium cervical and lumbar spinal cord data, differential expression was performed with DESeq2 (v1.50.2; Love, Huber and Anders, 2014) and counts were normalised with its median-of-ratios method. Significance was assessed with the Wald test, and log2 fold changes were shrunk with apeglm (Zhu, Ibrahim and Love, 2019). In KCL Brain Bank, the model was adjusted for age at death, sex, RNA integrity number and one surrogate variable. For Answer ALS data, the model was adjusted for sex, batch and surrogate variables and for ALS Consortium cervical and lumbar spinal cord, the model was adjusted for age at death, sex, RIN, tissue collection site and one surrogate variable. For this analysis, ALS Consortium motor cortex samples from medial, lateral and unspecified motor cortex were analysed together. Because donors contributed more than one sample, we used dream (variancePartition v1.40.2; Hoffman and Roussos, 2021) to run the analysis, following Humphrey et al. (2026). Counts were TMM-normalised (edgeR v4.8.2; Robinson and Oshlack, 2010) and transformed with voomWithDreamWeights (Law *et al*., 2014). The model included age at death, sex, RIN, collection site and two surrogate variables as fixed effects, with donor and motor cortex subregion as random effects. Degrees of freedom were estimated with the Satterthwaite approximation, followed by empirical Bayes moderation (limma v3.66.0).

Each analysis was run with cell type estimates added as covariates. For post-mortem tissue these were deconvolved cell type proportions and the most abundant cell type was omitted as the reference. For Answer ALS they were immunostaining percentages for SMI32, ISL1, NKX6.1, TUJ1, S100β and Nestin. All cell type covariates were z-scored, and in every dataset, P values were corrected with the Benjamini-Hochberg method. Genes with FDR < 0.05 were considered differentially expressed. Analyses were run in R v4.5.1.

### Gene set enrichment analysis

Gene set enrichment analysis (GSEA) was performed in R using fgsea (1.36.2) (Korotkevich *et al*., 2016) to test whether MitoCarta3.0 mitochondrial pathways (Rath *et al*., 2021) were enriched among genes differentially expressed between pwALS and controls. Each dataset was analysed separately. Ensembl gene identifiers were mapped to HGNC gene symbols using org.Hs.eg.db (3.22.0), so that all datasets were ranked on the same identifiers. Genes without a symbol or test result were removed.

Where several entries mapped to one symbol, the entry with the highest mean expression (DESeq2 baseMean or dream average log2 expression) was kept. Genes were ranked by the unshrunken test statistic from each differential expression model, the Wald statistic for DESeq2, and for dream the signed z-score equivalent of each gene’s p-value (z.std), because dream t-statistics have gene-specific degrees of freedom and are not directly comparable across genes. Two rankings were analysed for each dataset. In the genome-wide analysis, all tested genes were ranked, so that each pathway was compared against the whole transcriptome. In the mitochondrial-background analysis, the ranking was restricted to MitoCarta3.0 genes, so that each pathway was compared against other mitochondrial genes. This separates pathway-specific effects from a general shift in mitochondrial gene expression. Pathways from all levels of the MitoCarta3.0 hierarchy were tested, restricted to those with 5-2,000 genes present in the ranked list. Enrichment p-values were estimated with fgsea’s adaptive multilevel Monte Carlo procedure (eps = 0, random seed 42). P-values were adjusted for multiple testing within each dataset, model and background using the Benjamini-Hochberg method, and pathways with an adjusted p-value (FDR) < 0.05 were considered significantly enriched. Because pathways at different levels of the hierarchy share genes, these tests are not independent. Leading-edge genes were extracted for each pathway.

### Gene set variation analysis

Genes with >= 10 counts in at least as many samples as the smaller diagnostic group were retained (17,243-19,095 genes per tissue). Counts were transformed with DESeq2’s variance-stabilising transformation and technical variation was removed with limma::removeBatchEffect, with diagnosis protected in the design matrix. We regressed out sex, RIN, cell type proportion estimates and surrogate variables for KCL Brain Bank, sex, RIN, cell type proportion estimates, collection site and surrogate variables for ALS Consortium and sex, batch, immunostaining percentages and surrogate variables for Answer ALS. We scored the 149 MitoCarta3.0 pathways with GSVA (Hänzelmann, Castelo and Guinney, 2013; Fang, Liu and Peltz, 2023) if at least 5 member genes were expressed, resulting in 115 remaining pathways (114 in Answer ALS). Between 988 and 998 of the 1,035 MitoCarta pathway genes were detected per tissue. For this analysis ALS Consortium, one sample from each motor cortex donor was retained.

### Clustering

Samples were clustered on the GSVA scores of 36 tier 2 MitoCarta3.0 pathways. We used Ward’s hierarchical clustering (ward.D2) on Euclidean distances. The number of clusters, k, was chosen from 2 to 6 by maximising mean silhouette width. Stability was assessed as the adjusted Rand index between the hierarchical clusters and k-means clusters (50 random starts). Clusters were ordered by mean pathway score and labelled from low to high activity.

For single-pathway analyses, samples were labelled high or low by a two-component Gaussian mixture model (mclust). The model was used when its smaller component held at least 10% of samples. Otherwise, the top and bottom tertiles were compared and the middle tertile excluded.

We tested clusters for association with diagnosis, sex, age at death, RIN, post-mortem interval, collection site or batch, surrogate variables and cell-type estimates. Categorical variables were tested with Fisher’s exact test and continuous variables with Kruskal-Wallis tests (Benjamini-Hochberg (BH) correction within tissue). Cell-type estimates were deconvolution proportions for KCL Brain Bank and ALS Consortium and immunostaining percentages for Answer ALS. At the pathway level, each GSVA score was tested against each variable by Spearman correlation or Kruskal-Wallis test. BH correction was applied within each tissue for cluster tests, and within each variable and sample set for pathway tests.

### Exploratory survival analysis

Survival analyses were restricted to ALS cases. For KCL Brian Bank, the survival outcome was time to death in years. For ALS Consortium, it was disease duration (symptom onset to death) converted from months to years. All donors were deceased, so every case was an event. Individuals with missing time to event data were excluded.

We compared survival between high and low activity clusters with Kaplan-Meier curves and log-rank tests. Cox models estimated hazard ratios (HR) for clusters, with a likelihood-ratio test for cluster and a trend test across ordered clusters. Mean activity was modelled as a continuous score (HR per 1 SD). For each of the 36 pathways we estimated HRs for high vs low groups and per SD of score, with BH correction across pathways. Models were fitted unadjusted and adjusted for sex and age at onset.

Proportional hazards were checked with Schoenfeld residuals (cox.zph). Groups with fewer than 5 donors were excluded.

### Mito-nuclear co-expression analysis

Within each tissue tested, genes were retained if they had at least 20 counts in total and non-zero counts in every sample, leaving between 15,305 and 15,638 genes per dataset. Counts were transformed with a DESeq2 variance-stabilising transformation, blinded to diagnostic group. Following previously established methodology (Fairbrother-Browne *et al*., 2021), values more than 3 interquartile ranges beyond each gene’s quartiles were masked. Covariate effects were then removed from each gene by least-squares regression (limma removeBatchEffect), fitted on the unmasked values, with diagnostic group included as a protected term so that only covariate effects were removed and disease-associated variation was retained. For KCL Brain Bank, covariates were sex, age at death, RIN and one surrogate variable. For ALS Consortium tissues, they were sex, age at death, RIN and site of specimen collection, with two surrogate variables for motor cortex and one for the cervical and lumbar spinal cord. For Answer ALS, they were sex, batch and three surrogate variables. Cell type proportion estimates were also included as covariates. Where estimated proportions summed to one, the most abundant cell type was omitted as the reference. For Answer ALS, immunostaining percentages were included. Numeric technical covariates were mean-centred, cell type covariates were z-scored, and categorical covariates were treatment-coded. For this analysis ALS Consortium, one sample from each motor cortex donor was retained.

To measure nuclear-mitochondrial co-expression, Spearman’s ρ between each of the 13 protein-coding mtDNA genes and every nuclear-encoded gene were calculated for each dataset and diagnosis group separately, using the covariate-adjusted expression values and pairwise-complete samples. This gave one 13 x 15,292-15,625 correlation matrix per group per dataset.

### Significance testing of top level MitoCarta3.0 pathway gene set tests

Adapting the random gene set approach of Fairbrother-Browne et al. (2021), the test statistic for a pathway in one diagnosis group was the median of all its nuclear encoded pathway genes x mtDNA gene correlations. We compared this with 10,000 random sets of the same size drawn without replacement from expressed non-pathway genes. The shift was the observed median minus the mean of the 10,000 random-set medians.

To compare groups, we took the case-control difference in shift (shift in controls minus shift in ALS). The same random sets were used for both groups, so each random set also gave a null value of the difference. This paired null gave the p-value and a permutation-inverted 95% confidence interval. P-values were adjusted with the BH method across the 35 pathway x dataset tests, separately for each null and each comparison.

To ensure that case-control differences did not arise from an imbalance in diagnosis group size, ALS cases were subsampled without replacement to the number of controls in each dataset This was repeated 50 times, and correlations were recomputed for each draw and the was test re-run with the same random sets.

To make sure that a mismatch between highly expressed mitochondrial pathway genes and low expressed null genes were not driving observed shifts, we additionally repeated every gene set test with random sets matched on expression. Genes were binned into deciles of mean VST expression, and each random set drew as many genes from each decile as the pathway contained.

A case-control difference was called robust if it had P_BH_ < 0.05 under the expression-matched null, and P_unadj_ < 0.05 in at least 80% of subsampled draws. It was called power-limited if it was significant under both nulls but in fewer than 80% of draws, and unmatched-null only if it was significant only without expression matching.

## Results

### Mitochondrial translation genes are modestly enriched for ALS genetic risk

To test whether genetic variation in nuclear-encoded mitochondrial genes is enriched for ALS association, we performed MAGMA competitive gene-set analysis across tier 1 MitoCarta3.0 functional pathways (Figure 1, Table S1). Only mitochondrial central dogma showed significant enrichment after correction (β = 0.136, P_adj_ = 0.044), suggesting that genes in this category are more strongly associated with ALS than genes outside it.

**Figure 1.**
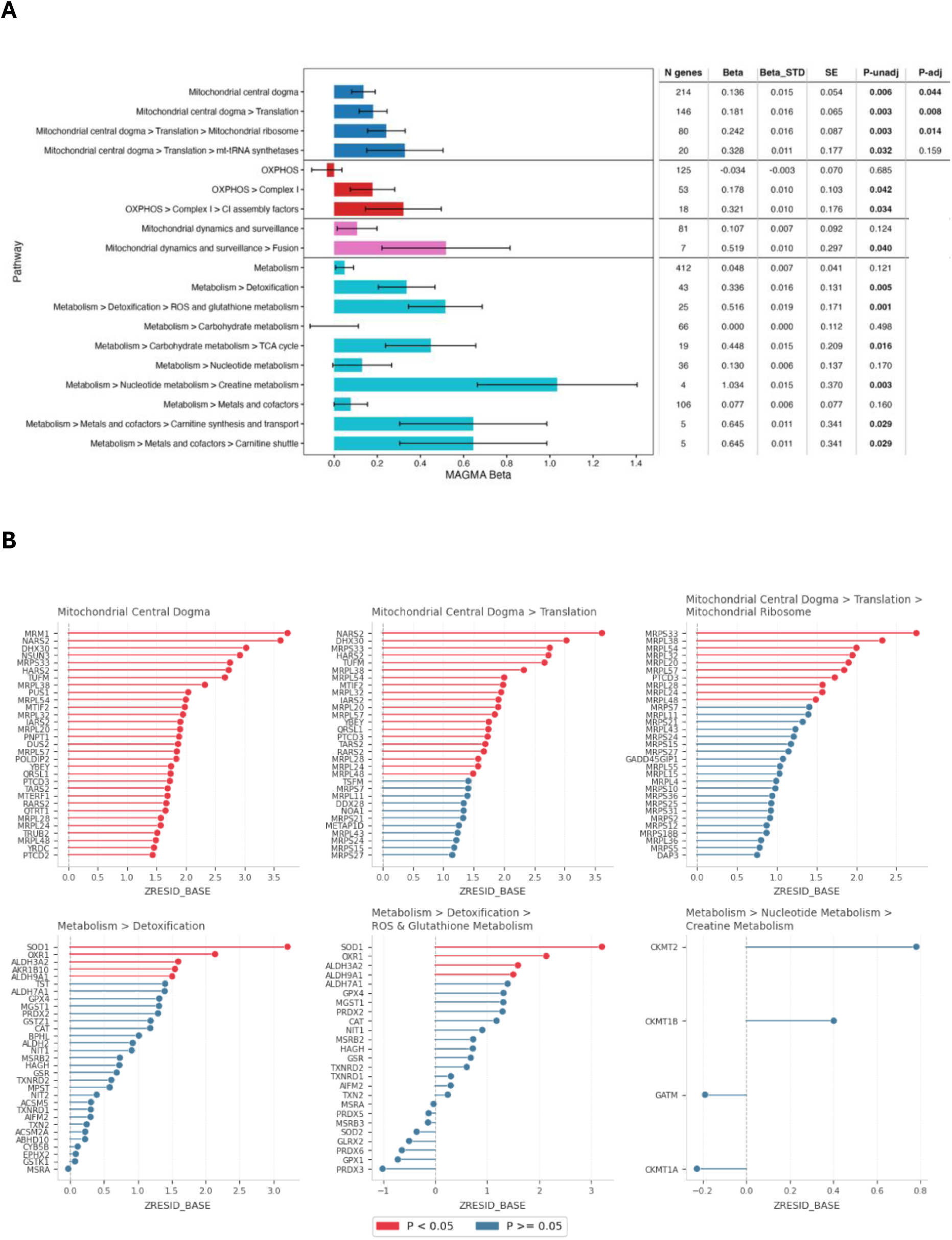
MAGMA gene-set analysis of mitochondrial pathways in ALS GWAS (Van Rheenen *et al*., 2021). **(A)** Competitive gene-set analysis of MitoCarta3.0 pathways using MAGMA. Top-level categories are shown with nominally significant sub-pathways and their parents (sublevels indicated by “>”). Bars show the MAGMA regression coefficient (β) ± SE, coloured by top-level category. The number of genes per pathway (N genes), β, standardised β (Beta_STD), SE, P_unadj_ and P_adj_ (hierarchical testing, see methods) are shown in the table to the right. **(B)** Gene-level results for pathways within the central dogma and metabolism gene sets. Genes are ranked by their residual MAGMA Z-statistic after adjustment for gene size and density (ZRESID_BASE). The top 31 genes are shown. Red, gene-level P_unadj_ < 0.05 and blue, P_unadj_ ≥ 0.05.

To identify the sub-processes driving this signal, and to detect enrichment that might be diluted within broader categories, we extended the analysis to tier 2 and tier 3 sub-pathways (Figure 1, Table S2-S3). Within the mitochondrial central dogma, enrichment localised to translation (β = 0.181, P_adj_ = 0.008) and, more specifically, to the mitochondrial ribosome (β = 0.242, P_adj_ = 0.014).

Sub-pathway analysis also revealed enrichment within metabolism. As metabolism was not significant in itself as a tier 1 pathway, the following results are considered as nominal. Within tier 2, detoxification showed some evidence of enrichment (β = 0.336, P_unadj_ = 0.005), with the strongest signal in its ROS and glutathione metabolism sub-pathway (β = 0.516, P_unadj_ = 0.001). Creatine metabolism, a tier 3 pathway within metabolism, also showed nominal evidence of enrichment (β = 1.034, P_unadj_ = 0.003), although this gene set contains only four genes. Finally, the metabolism tier 3 pathways: TCA cycle (β = 0.448, P_unadj_ = 0.016), Carnitine synthesis, and transport and Carnitine shuttle (both β = 0.645, P_unadj_ = 0.029) also showed a nominal signal. Similarly, within OXPHOS tier 2 pathways: complex I (β = 0.178, P_unadj_ = 0.042) and its CI assembly factors (β = 0.321, P_unadj_ = 0.034) and the tier 2 mitochondrial dynamics and surveillance pathway: fusion (β = 0.519, P = 0.040) were noted.

Next, to identify the genes contributing to these gene-set associations, we examined MAGMA gene-level results for the significantly enriched central dogma gene sets and also the nominally identified metabolism gene sets (Figure 1, Table S4-S6). Genes are reported descriptively at P_unadj_ < 0.05 without correction for multiple testing.

Within mitochondrial central dogma (214 genes), 31 genes showed nominal gene-level association with ALS risk. The strongest associations were with *MRM1* (Z = 3.95, P_unadj_ = 3.9 x 10⁻⁵) and *NARS2* (Z = 3.83, P_unadj_ = 6.4 x 10⁻⁵), followed by *DHX30*, *NSUN3*, *MRPS33*, *HARS2* and TUFM (all P_unadj_ < 0.002). 20 of the 31 genes belong to translation, including five mitochondrial tRNA synthetases (*NARS2*, *HARS2*, *IARS2*, *TARS2*, *RARS2*), the translation factors *TUFM* and *MTIF2*, and ten mitoribosomal subunits. Nominally significant mitoribosomal genes spanned both the large and small subunits: *MRPS33* (Z = 2.97, P_unadj_ = 0.0015) and *MRPL38* (Z = 2.55, P_unadj_ = 0.0055), followed by *MRPL54, MRPL32*, *MRPL20*, *MRPL57*, *PTCD3*, *MRPL28*, *MRPL24* and *MRPL48.* Most of the remaining 11 are rRNA and tRNA modification enzymes (*MRM1*, *NSUN3*, *PUS1*, *DUS2*, *ǪTRT1*, *TRUB2*, *YRDC*), alongside *PNPT1*, *PTCD2*, *MTERF1* and *POLDIP2*. Gene-level signal therefore spans the mitochondrial RNA processing and translation pathway, from RNA modification through to the mitoribosome itself.

Within metabolism, the strongest gene-level association was with *SOD1* (Z = 3.43, P_unadj_ = 3.0 x 10⁻⁴). *SOD1* was the top gene in both the detoxification sub-pathway (43 genes; five nominal genes) and its ROS and glutathione metabolism component (25 genes; four nominal genes). The other nominal genes in this component were *OXR1* (Z = 2.36, P_unadj_ = 0.009), *ALDH3A2* (P_unadj_ = 0.035) and *ALDHSA1* (P_unadj_ = 0.042). None of the 4 genes in the Creatine metabolism sub-pathway reached nominal significance.

Overall, beyond recovering the known ALS association with *SOD1*, gene-level results implicate mitochondrial RNA processing and translation as a source of ALS genetic risk. This signal is distributed broadly across the pathway and is most specific to the mitoribosome.

### Expression based annotation of MAGMA prioritised genes

To understand the directionality of the genes driving MAGMA gene-set associations, we examined TWAS and SMR results from (Pain *et al*., 2024) for the 33 of the 36 genes identified in the MAGMA analysis (Figure 2, Tables S7-S8), where data were available. As TWAS and SMR used the same GWAS as the MAGMA analysis, and genes were selected on their MAGMA signal, these results annotate the direction of effect rather than independently replicating the association. To separate associations likely to reflect a shared causal variant from those driven by linkage between distinct eQTL and GWAS variants, we highlight two kinds of result. For TWAS, we report the ratio PP4/(PP3+PP4), which indicates whether a shared causal variant is favoured over distinct linked variants, given that both traits are associated. Because this ratio can be high even when the absolute posterior probability of colocalisation (PP4) is low, we report both. For SMR, they are nominal associations that passed the HEIDI test (P_HEIDI > 0.05). Of the 36 MAGMA prioritised genes, 33 had at least one expression model, 20 showed at least one nominal association (P_unadj_ < 0.05) and 4 reached P_FDR_ < 0.05 in the original analysis (*NARS2*, *MRPS33*, *MRPL28* and *DUS2*), all within the mitochondrial central dogma.

**Figure 2.**
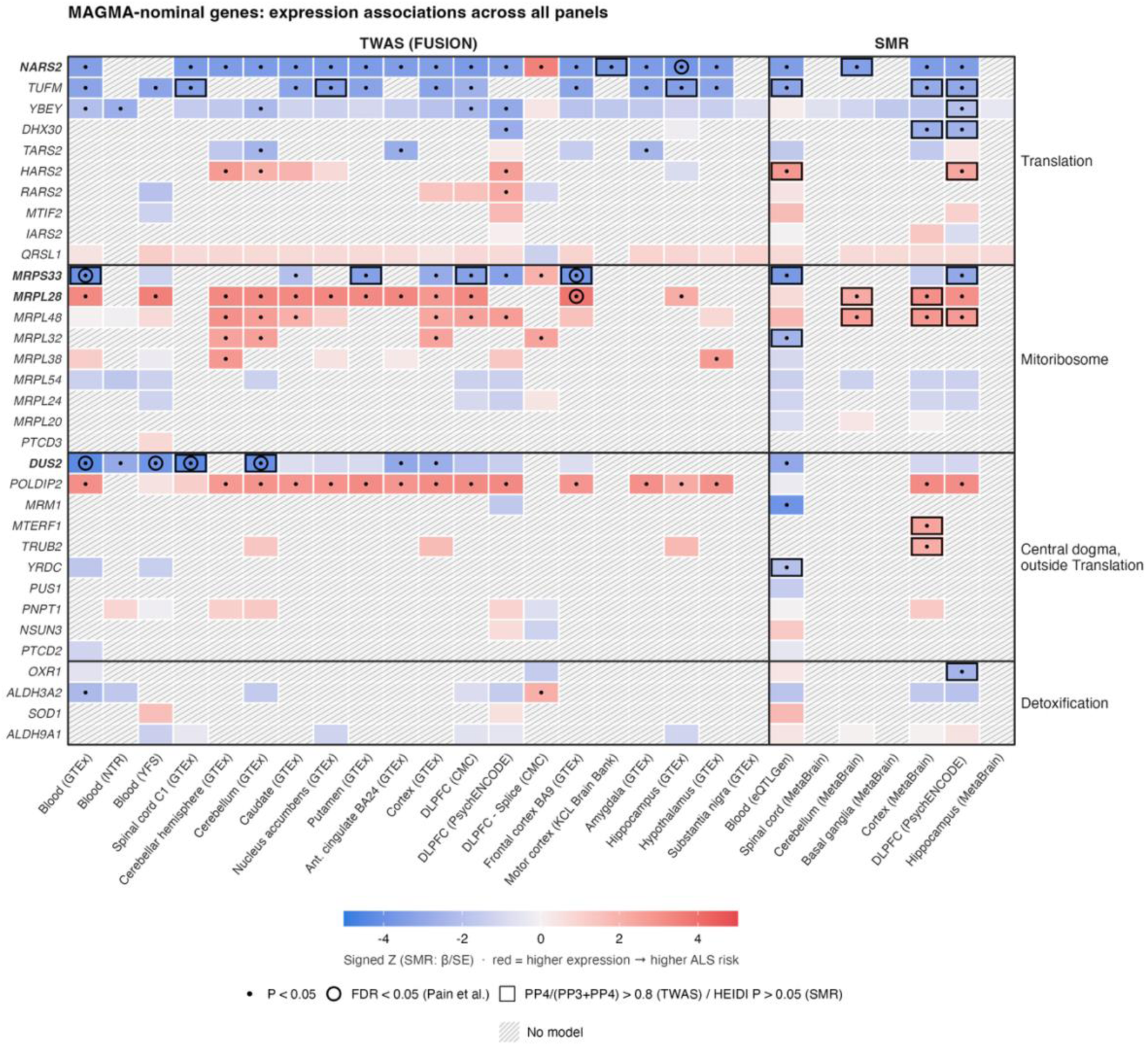
Expression associations of MAGMA-nominal mitochondrial genes with ALS risk. TWAS (FUSION) and SMR results from Pain et al. (2024) are shown for the 33 of 36 MAGMA-nominal genes with at least one expression model. *ǪTRT1*, *MRPL57* and *AKR1B10* had no model and are not shown. Rows are grouped by MitoCarta pathway (translation excluding the mitoribosome; mitoribosome; central dogma genes outside translation; detoxification) and ordered within each group by strength of expression support. Columns show the 20 FUSION panels (left) and 7 SMR panels (right), grouped by tissue. Cell colour shows the signed Z-statistic (FUSION TWAS Z; SMR β/SE). Red indicates that higher genetically predicted expression is associated with increased ALS risk, blue that lower expression is. Dots mark P_unadj_ < 0.05, rings mark P_FDR_ < 0.05 as reported by Pain et al. Squares mark TWAS associations with colocalisation support (PP4/(PP3 + PP4) > 0.8) or SMR associations that passed the HEIDI test (P_HEIDI > 0.05), indicating that the expression and ALS associations are more likely to share a causal variant rather than to reflect linkage between distinct variants. Hatched cells indicate no expression model for that gene in that panel. Genes in bold reached P_FDR_< 0.05 in at least one panel. The DLPFC-Splice (CMC) panel models splicing rather than total expression, so its direction is not directly comparable with the expression panels. DLPFC, dorsolateral prefrontal cortex; BA, Brodmann area; GTEx, Genotype-Tissue Expression; CMC, CommonMind Consortium; NTR, Netherlands Twin Register; YFS, Young Finns Study; KCL, King’s College London.

Among mitoribosomal genes, lower genetically predicted expression of *MRPS33* was linked to increased ALS risk. The association reached P_FDR_ < 0.05 in blood (GTEx) and brain (GTEx Frontal cortex BA9) TWAS panels. Where colocalisation was tested, the PP4/(PP3+PP4) ratio favoured a shared variant over distinct linked variants (0.92–0.96 in three brain panels), but absolute PP4 was low (0.04–0.34), so evidence that the expression and ALS signals share a causal variant is weak. This direction was consistent across 8 of 9 TWAS panels and 3 SMR panels, with both nominal SMR associations, blood (eQTLGen) and DLPFC (PsychENCODE) passing the HEIDI test. In contrast, higher predicted expression of *MRPL28* was associated with increased ALS risk. This directionality was consistent across 12 TWAS panels and 4 SMR panels, and the association reached P_FDR_ < 0.05 in brain (GTEx Frontal cortex BA9),but did not meet the colocalisation threshold in any of the three panels tested. *MRPL48* and *MRPL38* showed a similar positive pattern.

Within translation, *NARS2* was nominally associated in 17 TWAS panels and 4 SMR panels. Lower expression was linked to increased ALS risk in every panel except the DLPFC splicing panel, reaching P_FDR_ < 0.05 in hippocampus (GTEx), although colocalisation support in that panel was below threshold.

Outside of translation pathways, lower predicted expression of *DUS2*, a tRNA dihydrouridine synthase, was likewise associated with increased ALS risk. Two of its four FDR significant TWAS panels showed colocalisation support with PP4 of 0.85 (cervical spinal cord, GTEx) and 0.72 (cerebellum, GTEx). The direction of effect was the same in all 16 panels with a model.

Expression evidence was therefore concentrated in the mitochondrial central dogma, particularly translation, but its direction was gene-specific rather than uniform across the pathway. For example, lower predicted expression of *MRPS33* was associated with increased ALS risk, whereas higher expression of *MRPL28* was associated with increased ALS risk, despite the two genes encoding mitochondrial proteins. *MRM1* and *NSUN3*, the two strongest MAGMA genes, had expression models in few panels and showed no robust expression support, suggesting that their MAGMA signal may act through mechanisms not captured by bulk-tissue eQTLs. Detoxification genes showed no consistent expression association, including *SOD1*.

For the 33 genes identified though MAGMA analysis with corresponding TWAS and SMR models, we also asked if they were differentially expressed in ALS tissue. To do this, we compared the strongest direction of effect from TWAS and SMR (Pain *et al*., 2024) with the direction of effect from differential gene expression (DGE) in five datasets: post-mortem motor cortex from KCL Brain Bank and ALS Consortium, post-mortem cervical and lumbar spinal cord form ALS Consortium, and iPSC-derived motor neurons from Answer ALS (Figure 3).

**Figure 3.**
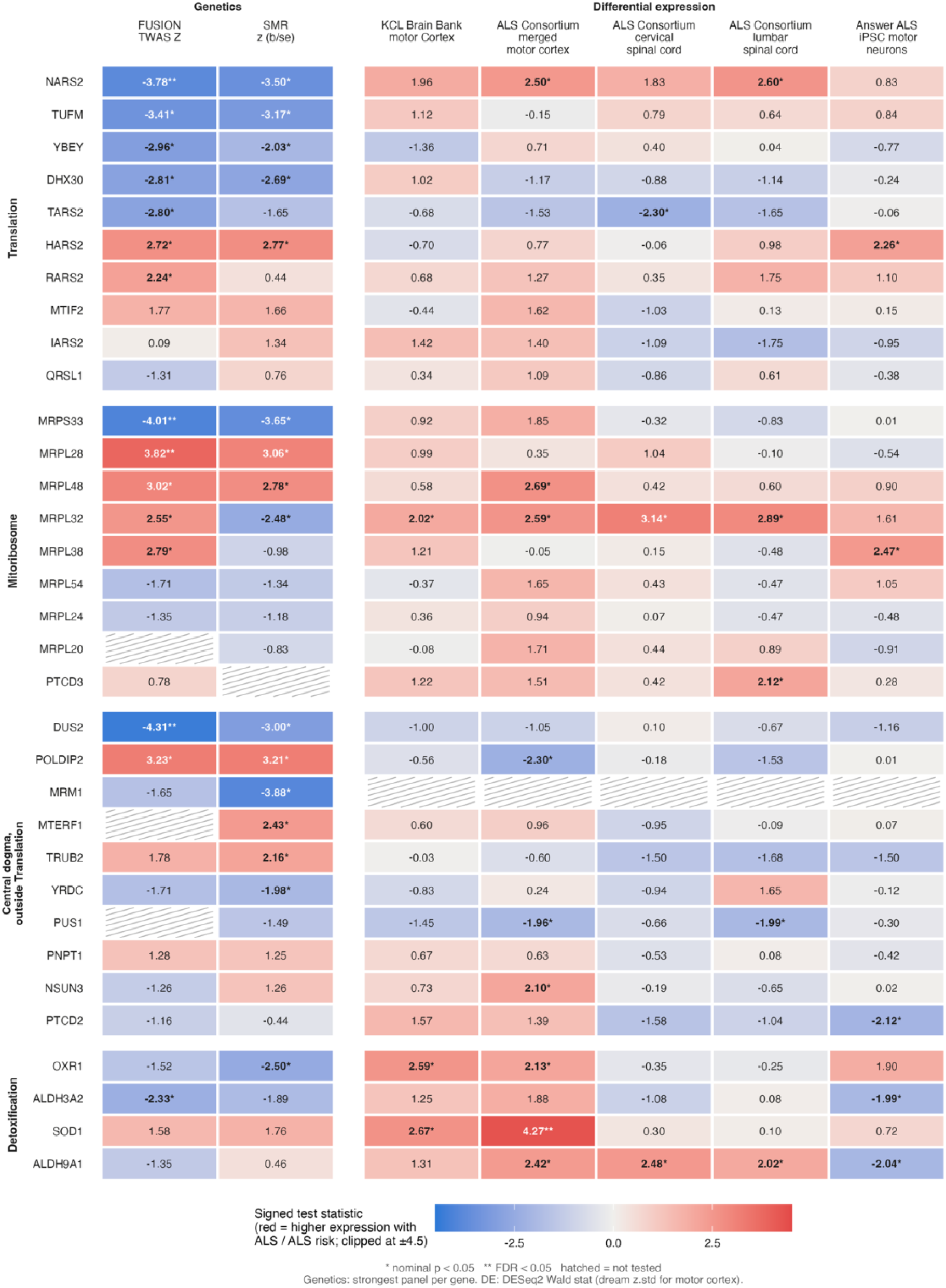
Genetically predicted expression and differential expression in ALS for selected mitochondrial genes. Heatmap shows signed test statistics for 33 mitochondrial genes (rows). The left panel shows genetic associations between gene expression and ALS risk and for each gene, the result from the expression panel (TWAS or SMR) with the lowest p-value is shown. The right panel shows differential expression between ALS cases and controls in five datasets, adjusted for estimated cell-type proportions. Values are DESeq2 Wald statistics, except for motor cortex, which shows the standardised moderated statistic from dream (variancePartition). Red indicates higher genetically predicted expression associated with ALS risk (left panel) or higher expression in ALS cases than in controls (right panel) and blue indicates the opposite. The colour scale is truncated at ±4.5. * nominal p < 0.05; ** false discovery rate (FDR) < 0.05 (Benjamini-Hochberg, applied across all genes tested within each method or dataset). Hatched cells mark genes that were not tested: no FUSION model (*MRPL20*, *MTERF1*, *PUS1*), no SMR result (*PTCD3*), or not present in the differential expression results (*MRM1*).

Among the 33 prioritised genes, the four that passed TWAS FDR correction (*DUS2*, *MRPS33*, *MRPL28* and *NARS2*) were not significantly differentially expressed in any dataset. For NARS2, lower predicted expression was associated with ALS risk (Z = −3.78), but expression was nominally higher in ALS motor cortex and lumbar spinal cord (DE statistics 2.50 and 2.60). Only one of the 33 genes was differentially expressed after FDR correction: *SOD1* was higher in ALS motor cortex (padj = 0.004), and its genetic associations, although not significant, pointed the same way (TWAS Z = 1.58; SMR z = 1.76). *MRPL32* showed the most consistent DE, being nominally higher in ALS in four of the five datasets, but its TWAS and SMR estimates pointed in opposite directions.

To test this more systematically, we extended the comparison to all MitoCarta3.0 genes with genetic and DGE results (Table S9). In every dataset, and for both TWAS and SMR, genetic and DGE effects agreed in direction for 48-52% of genes, no more often than expected by chance (all binomial p > 0.26). Genetic and DGE statistics were not correlated (Spearman |ρ| ≤ 0.06, all p > 0.06) and concordance did not increase when we restricted the analysis to genes with nominal genetic associations (39-55%), or to genes nominally significant in both analyses (27-75%). Together, these results suggest that the mitochondrial expression changes seen in ALS tissue are largely separate from the genetically determined expression effects linked to ALS risk.

### Mitochondrial gene sets are most strongly altered in ALS motor cortex

To understand if mitochondrial pathways are altered as a whole in ALS, we performed gene set enrichment analysis (GSEA) of MitoCarta3.0 pathways in five ALS versus control differential expression datasets. Each analysis was run twice. A genome-wide background tested whether mitochondrial pathways were enriched relative to all expressed genes (Figure 4, Table S10), and a background restricted to MitoCarta3.0 genes tested whether particular mitochondrial pathways stood out from mitochondrial genes as a whole (Figure S1, Table S11).

**Figure 4.**
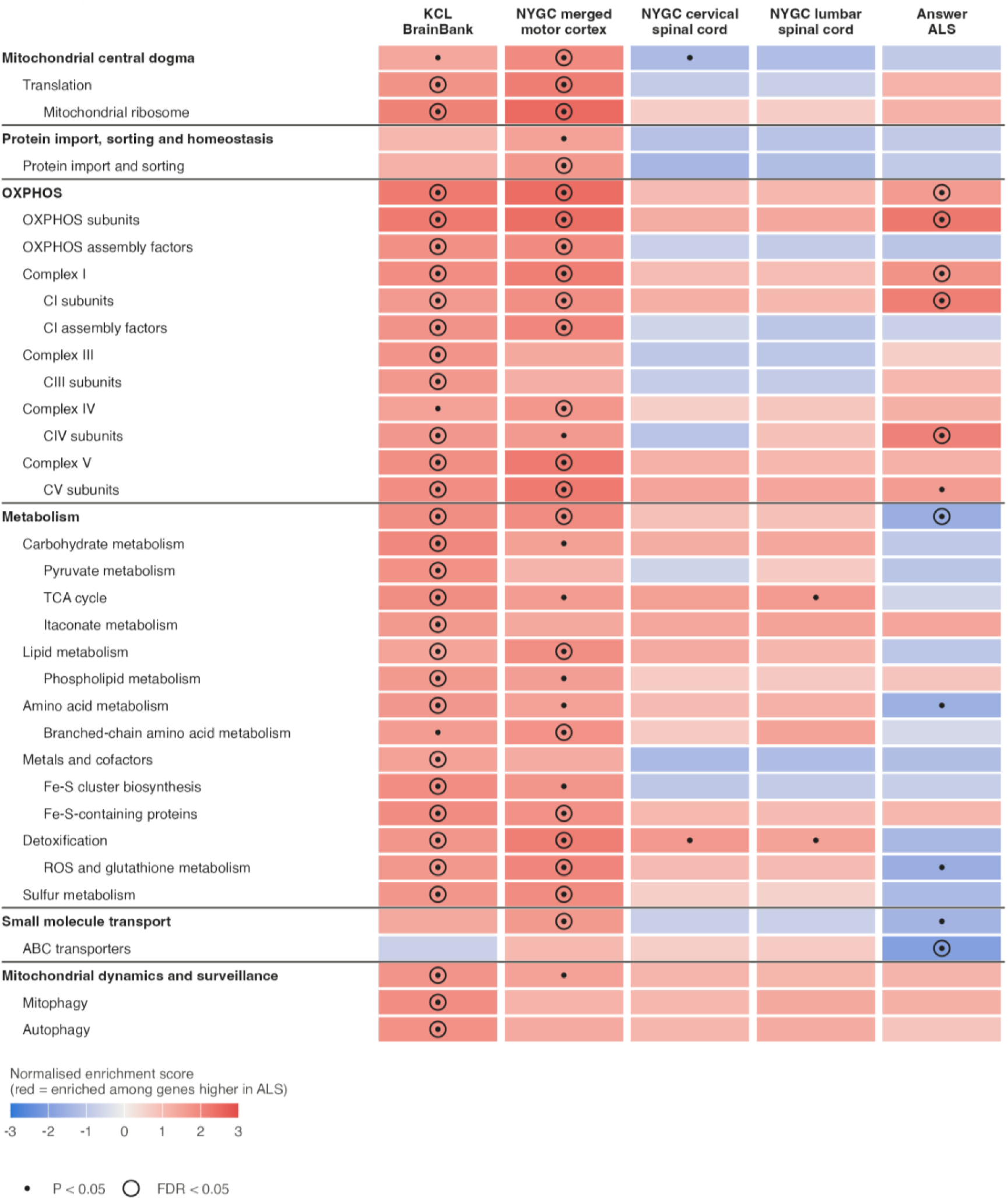
Mitochondrial pathway enrichment in ALS tissue and iPSC-derived motor neurons. Gene set enrichment analysis (fgsea) of MitoCarta3.0 mitochondrial pathways in ALS compared with controls. Columns are the five datasets: KCL Brain Bank motor cortex, NYGC motor cortex (merged), NYGC cervical spinal cord, NYGC lumbar spinal cord and Answer ALS iPSC-derived motor neurons. Rows are pathways with P_FDR_ < 0.05 in at least one dataset, together with their parent pathways. Pathways are grouped by MitoCarta top-level category, with indentation showing hierarchy depth. Genes were ranked by the DESeq2 Wald statistic (KCL BrainBank, ALS Consortium spinal cord, Answer ALS) or the dream moderated z-statistic (ALS Consortium motor cortex), from models adjusted for cell-type composition. Enrichment was tested against a genome-wide background. Cell colour shows the normalised enrichment score (NES): red indicates enrichment among genes more highly expressed in ALS, and blue indicates enrichment among genes more lowly expressed in ALS. A dot marks nominal P < 0.05, and a circled dot marks P_FDR_ < 0.05 (Benjamini-Hochberg, within each dataset).

Against the genome-wide background, enrichment was strongest and most widespread in motor cortex. Thirty pathways passed P_FDR_ < 0.05 in the KCL Brain Bank and 21 in ALS Consortium motor cortex, all of them higher in ALS. Overall, 88% and 91% of tested pathways had a positive NES. Most of this cortical enrichment was not pathway-specific. With the mitochondrial background, significant pathways dropped from 30 to none in the KCL Brain Bank and from 21 to four in ALS Consortium motor cortex. In ALS Consortium spinal cord and iPSC motor neurons, NES values were similar between the two analyses. The widespread enrichment in motor cortex therefore largely reflects higher expression of mitochondrial genes overall in ALS. This global shift was not seen in spinal cord or iPSC motor neurons.

Against the genome-wide background, OXPHOS was the top-ranked pathway in both cortex cohorts (KCL Brain Bank NES = 2.20, P_FDR_ = 1.0 x 10⁻⁸; ALS Consortium NES = 2.44, P_FDR_ = 2.2 x 10⁻¹⁰), and OXPHOS subunits were the top pathway in Answer ALS iPSC motor neurons (NES = 2.25, P_FDR_ = 2.7 x 10⁻⁶). With the mitochondrial background, OXPHOS subunits were enriched beyond the general mitochondrial shift in all five datasets, although the result passed FDR correction only in ALS Consortium motor cortex and Answer ALS iPSC motor neurons.

The mitochondrial ribosome was also enriched in both cortex cohorts against the genome-wide background (KCL Brain Bank NES = 2.07, P_FDR_ = 2.0 x 10⁻⁴; ALS Consortium NES = 2.48, P_FDR_ = 5.7 × 10⁻⁸). This held against the mitochondrial background in ALS Consortium motor cortex (NES = 1.72, P_FDR_ = 0.027) and at nominal significance in the KCL Brain Bank (NES = 1.42, P = 0.028).

In spinal cord, no pathway passed P_FDR_ < 0.05 against the genome-wide background. Against the mitochondrial background, however, cervical spinal cord showed significant depletion of mitochondrial central dogma (NES = −1.57) and mtRNA metabolism (NES = −1.74), and enrichment of detoxification (NES = 1.82; all P_FDR_ = 0.045). Lumbar spinal cord showed the same direction for these pathways at nominal significance only.

In Answer ALS iPSC motor neurons, results were very similar under both backgrounds, and the same seven pathways passed FDR. Mitochondrial metabolism as a whole was depleted (genome-wide NES = −1.58, P_FDR_ = 0.0019; mitochondrial NES = −1.61, FDR = 0.0014), as were ABC transporters (NES = −1.93, P_FDR_ = 0.037). Several metabolic subpathways enriched in motor cortex, including ROS and glutathione metabolism and amino acid metabolism, were nominally depleted (P < 0.05).

### Mitochondrial activity clusters are weakly defined, associated with diagnosis in motor cortex, and with survival only in lumbar spinal cord

In all 5 tissues tested, gene set variation analysis (GSVA) scores of mitochondrial pathway activity separated samples into two clusters, low and high (Figure 5). Cluster structure was modest, with mean silhouette widths ranging from 0.22 to 0.33 (Table 1). Agreement between hierarchical and k-means clusters was moderate in KCL Brain Bank, ALS Consortium motor cortex and cervical spinal cord, (adjusted Rand index 0.75-0.84) but poor in ALS Consortium lumbar spinal cord and Answer ALS iPSC motor neurons (0.32 and 0.59; Table 1).

**Figure 5.**
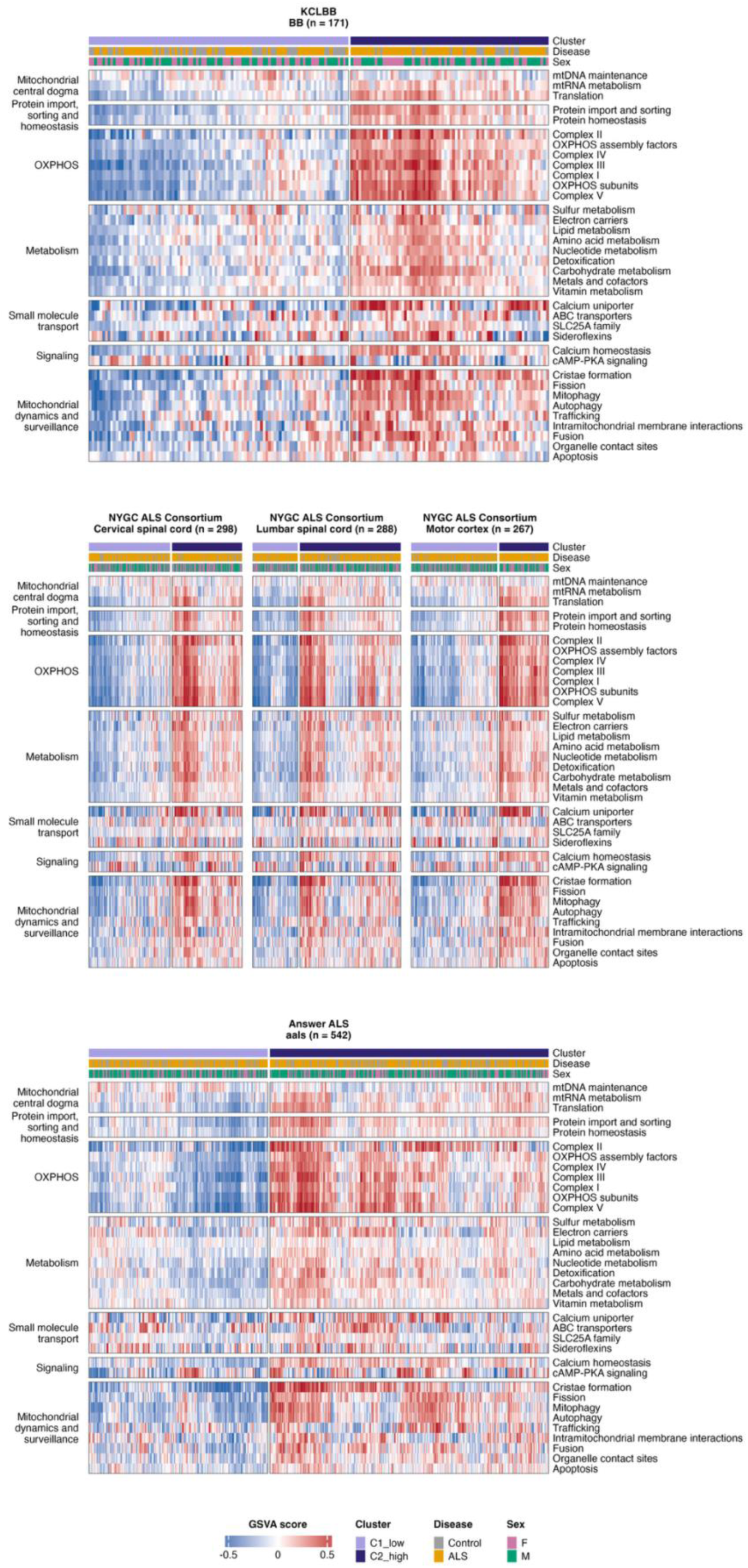
Mitochondrial pathway activity across ALS and control samples. Heatmaps of GSVA scores for the 36 tier 2 MitoCarta3.0 pathways (rows) in the KCL Brain Bank, the NYGC ALS Consortium (motor cortex, cervical spinal cord and lumbar spinal cord) and Answer ALS iPSC-derived motor neurons. Panel titles give the cohort, tissue and number of samples. Red indicates higher and blue lower pathway activity relative to other samples in the same tissue. All panels share one colour scale, capped at the 99th percentile of absolute scores. Pathways are grouped by their top-level MitoCarta3.0 category and ordered within each category by Ward hierarchical clustering of scores across all tissues, so rows are directly between panels. Samples (columns) are split into the pathway-activity clusters identified in each tissue. Ward hierarchical clustering with the number of clusters chosen by mean silhouette width, numbered from lowest (C1) to highest mean activity. Within each cluster, samples are also ordered by Ward hierarchical clustering. Annotation bars show cluster, diagnosis (ALS or control) and sex (F, female; M, male).

**Table 1.** Cluster structure and associations by tissue. ARI = adjusted Rand Index.

| **Cohort and Tissue** | **Chosen k** | **Mean silhouette** | **ARI vs k-means** | **C1_low n** | **C2_high n** | **Total n** | **Cluster vs disease p** | **Cluster vs disease BH** |
|---|---|---|---|---|---|---|---|---|
| KCL Brain Bank Motor Cortex | 2 | 0.285 | 0.842 | 97 | 74 | 171 | 0.0060 | 0.0332 |
| ALSC. Motor cortex | 2 | 0.328 | 0.773 | 170 | 97 | 267 | 0.0028 | 0.0447 |
| ALSC. Cervical spinal cord | 2 | 0.287 | 0.749 | 160 | 138 | 298 | 0.5370 | 0.9150 |
| ALSC. Lumbar spinal cord | 2 | 0.247 | 0.322 | 89 | 199 | 288 | 0.0913 | 0.7430 |
| iPSC motor neurons (aals) | 2 | 0.222 | 0.594 | 212 | 330 | 542 | 0.5570 | 0.8630 |

Cluster membership was associated with ALS diagnosis in both motor cortex cohorts (Fisher P_FDR_ < 0.05, Table 1), but was not associated with sex, age at death, RIN, post-mortem interval, collection site, surrogate variables, or cell type proportion estimates, except in the KCL Brain Bank (Table S12). In this dataset, cluster membership was associated with RIN and SV1.

We next carried out an exploratory examination of whether cluster membership was associated with survival using Cox proportional hazards regression, comparing the C2_high cluster with C1_low.

Mitochondrial activity clusters were associated with survival in only one of the four tissues (Figure 6). In ALS Consortium lumbar spinal cord (232 donors), donors in the high-activity cluster had longer disease duration than those in the low-activity cluster (HR 0.62, log-rank p = 0.0008; Table S13). The association was weaker but persisted after adjustment for sex and age at symptom onset (HR 0.69, p = 0.013). Mean mitochondrial activity analysed as a continuous score pointed the same way (HR 0.84 per SD, p = 0.020; Table S14), but it was no longer significant after adjustment for sex and age at symptom onset (HR 0.89, p = 0.14). Cluster structure in lumbar spinal cord was the weakest of all the datasets (mean silhouette 0.247, agreement between clustering methods ARI 0.32). This suggests mitochondrial activity in this tissue is closer to a continuum than two distinct groups, and the two-cluster split may not be stable. The continuous-score result is less affected by this, but is not significant after adjustment.

**Figure 6.**
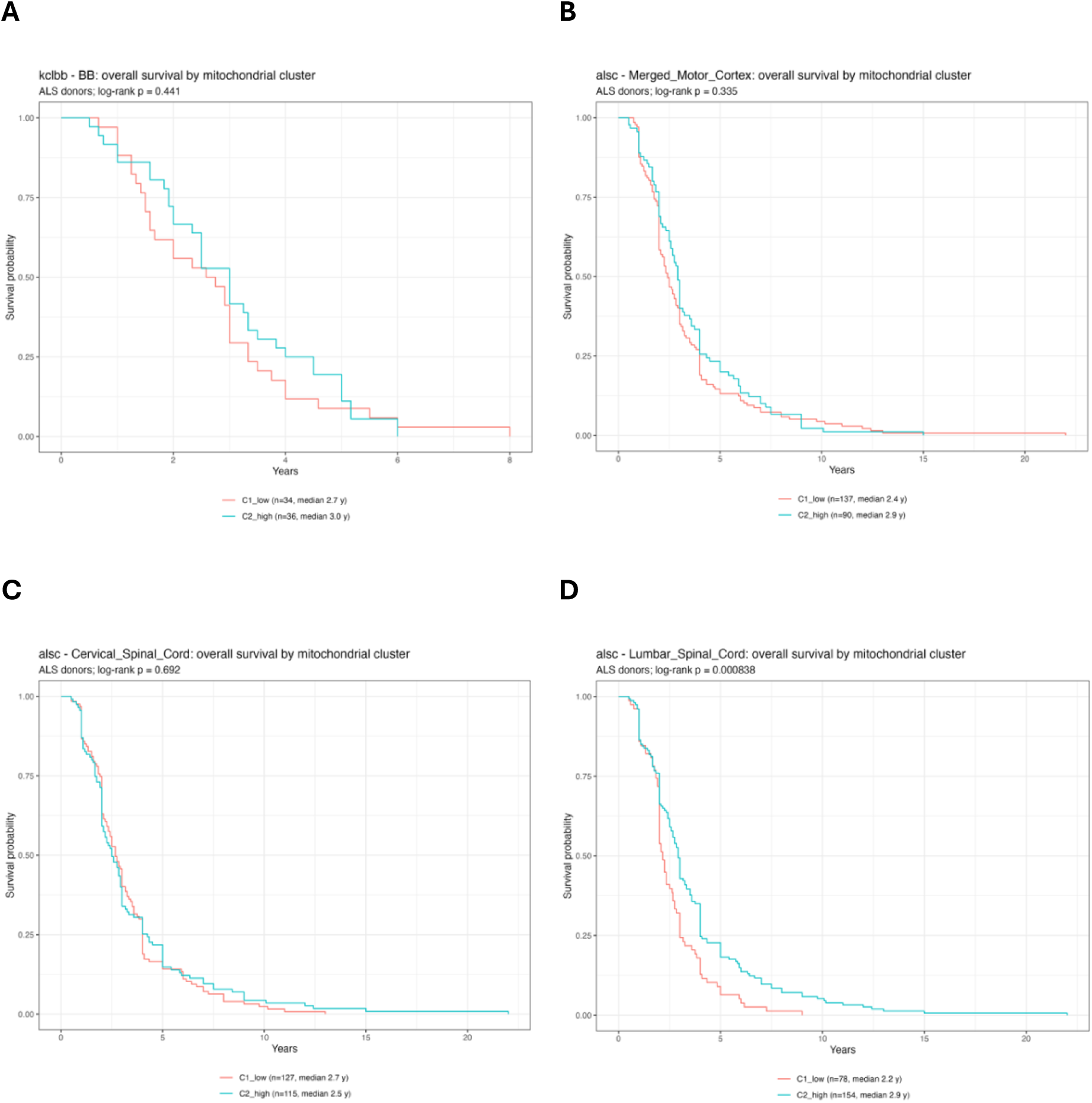
Overall survival by mitochondrial cluster in end stage ALS tissues. Kaplan-Meier curves of survival curves (unadjusted models) for ALS in **(A)** KCL Brain Bank, **(B)** ALS Consortium merged motor cortex, **(C)** ALS Consortium cervical spinal cord and **(D)** ALS Consortium lumbar spinal cord. All donors were deceased at the time of tissue collection, so no observations were censored.

Clusters were not associated with survival in cervical spinal cord (HR 0.95, p = 0.68), motor cortex (HR 0.88, p = 0.35) or the KCL brain bank samples (HR 0.85, p = 0.51; 70 donors). Continuous mean activity was also null in all three (HR 0.87 to 1.00 per SD).

The proportional hazards assumption was assessed using Schoenfeld residuals (cox.zph). There was no evidence of violation in any cluster model, unadjusted or adjusted (global test p ≥ 0.17 across all tissues; lumbar spinal cord p = 0.42 unadjusted and p = 0.53 adjusted; Table S13-S15). The only nominal departure among the continuous mean-activity models was in cervical spinal cord (unadjusted p = 0.036). That association was null, and the departure was not present after adjustment (p = 0.16).

### Mitochondrial pathway activity and survival in ALS

To determine whether specific mitochondrial functions were associated with survival, we tested scores for 36 mitochondrial tier 2 pathways in each cohort and tissue using Cox proportional hazards regression. Pathway scores were modelled as continuous variables (hazard ratio (HR) per standard deviation (SD)) and as high versus low groups, with and without adjustment for sex and age at onset. P-values were corrected for multiple testing within each tissue using the Benjamini-Hochberg method (Table S15).

No pathway was associated with survival after BH correction in cervical spinal cord or motor cortex (Figure 7). In lumbar spinal cord, 11 of 36 tier 2 pathways passed BH correction (< 0.05) on at least one unadjusted test, either high vs low group or per-SD score. Ten of these pointed towards longer survival with higher activity (HR about 0.79 to 0.87 per SD), including: mitochondrial translation, protein homeostasis, OXPHOS assembly factors, complex III, lipid, nucleotide and vitamin metabolism, detoxification, SLC25A carriers and fission. cAMP-PKA signalling was the one exception, where higher scores went with shorter survival (HR 1.20 per SD). Higher organelle contact site activity was nominally associated with longer survival in KCL Brain Bank motor cortex (adjusted HR 0.78 per SD) and ALS Consortium cervical spinal cord (adjusted HR 0.85 per SD), but did not survive multiple test correction. The adjusted models included fewer donors (215 vs 232) because age at onset was missing for some, so differences in results may partly reflect reduced sample size rather than confounding.

**Figure 7.**
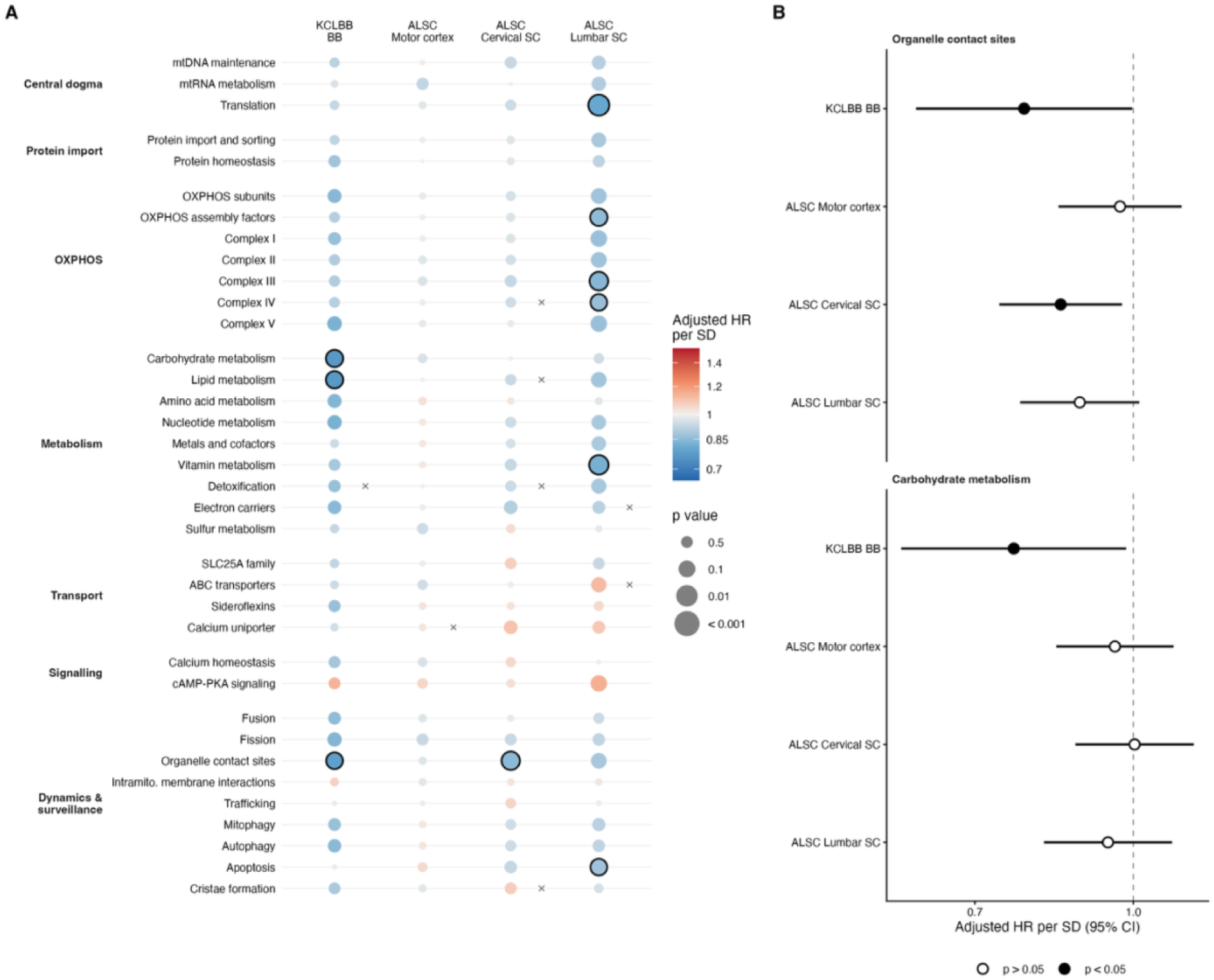
Association of mitochondrial pathway activity with survival in ALS. **(A)** Adjusted hazard ratios (HRs) for death per 1 standard deviation (SD) increase in GSVA score for 36 MitoCarta3.0 pathways (rows), grouped by functional category, in KCL Brain Bank and ALSC cervical spinal cord, lumbar spinal cord and motor cortex (columns). Point colour indicates the HR (blue, HR < 1, longer survival with higher pathway activity; red, HR > 1, shorter survival). Point size indicates the p-value and black outlines mark nominal significance (p < 0.05). No association remained significant after Benjamini–Hochberg correction within each tissue (all P_BH_ ≥ 0.10). Crosses (x) indicate violation of the proportional hazards assumption for the pathway term (Schoenfeld residual test, p < 0.05). **(B)** Adjusted HRs per SD (95% confidence intervals, log scale) for organelle contact sites and carbohydrate metabolism across tissues. Filled points indicate p < 0.05; the dashed line marks HR = 1.

Overall, the cell-type-adjusted analysis gives at most a tissue-specific, hypothesis-generating signal and tissue sampled earlier in the disease would be better for testing.

### Co-expression of mitochondrial pathway gene sets with mtDNA-encoded genes

As the nuclear encoded and mitochondria encoded mitochondrial genes are required to co-ordinate in order to function appropriately, we asked if the nuclear genes in top-level MitoCarta3.0 pathways showed altered co-expression with the 13 protein-coding mtDNA genes in ALS, after adjusting for technical covariates and cell type composition.

Of 35 pathway x dataset comparisons, 9 showed a robust case-control difference in nuclear-mitochondrial co-expression, 7 were significant but power-limited, 4 were significant only under the unmatched null, and 15 were not significant (Figure 8, Table S16). At least one robust difference was observed in every dataset. In the four post-mortem tissues, MitoCarta3.0 pathway genes were more negatively correlated with mtDNA genes, relative to random genes, in ALS than in controls.

**Figure 8.**
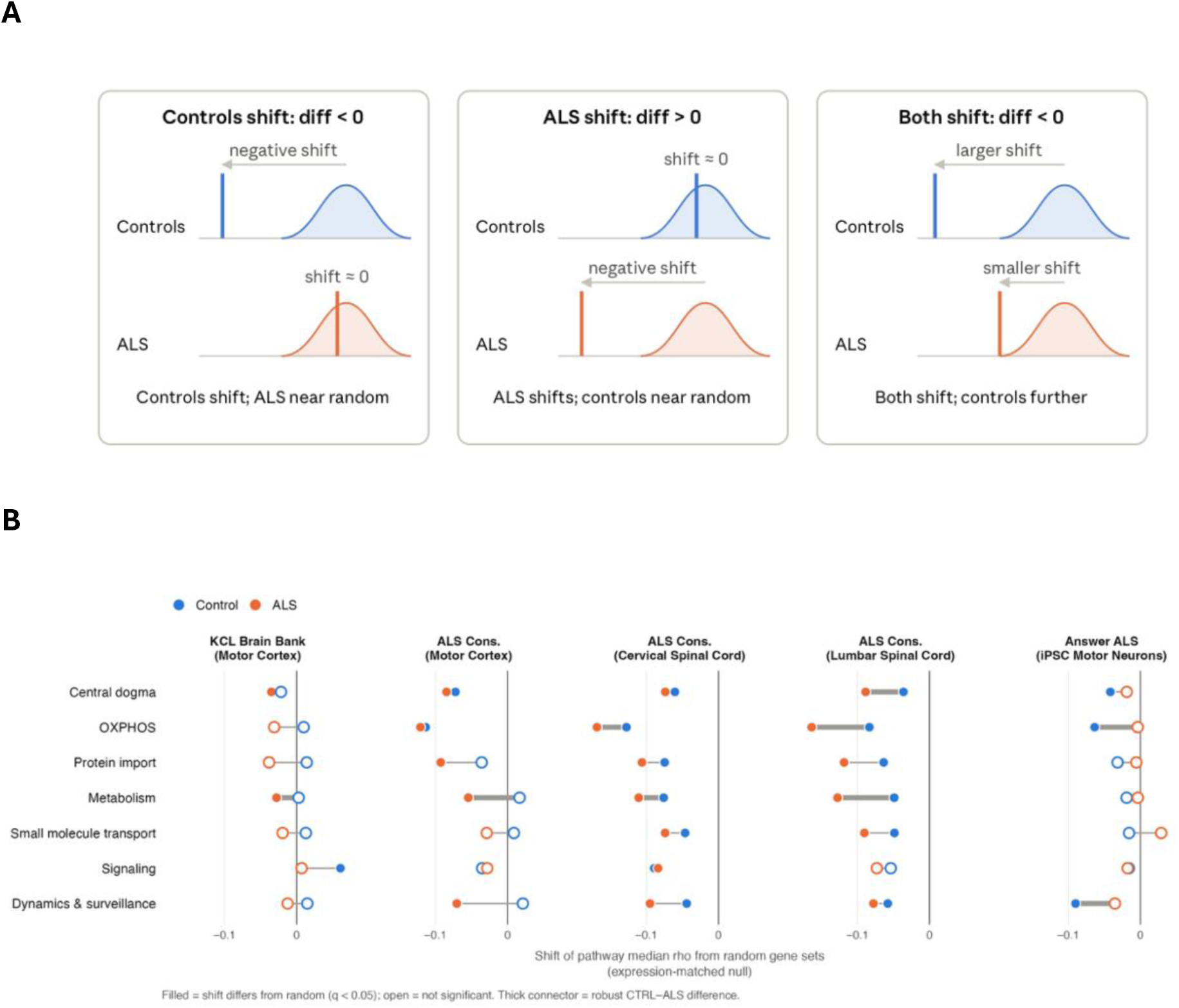
Co-expression of mitochondrial pathways with mtDNA-encoded genes differs between ALS and controls in a region-dependent manner. **(A)** How to read a “shift” and the case-control difference using toy data. Each curve shows the medians of 10,000 random gene sets. Each vertical line represents a pathway observed median. A shift describes the pathway observed median minus the curve centre. The difference measures the shift in controls minus the shift in ALS. **(B)** For each mitochondrial pathway (rows) and dataset (columns), the shift in co-expression with the 13 mtDNA-encoded protein-coding genes relative to random gene sets, in controls (blue) and ALS (orange). The shift is the median Spearman ρ across all pathway gene x mtDNA gene pairs minus the mean of the same statistic for 10,000 random gene sets of equal size. Random genes were drawn from expressed non-pathway genes and matched to the pathway on mean expression (deciles of mean VST expression). Filled circles, P_BH_ < 0.05 (two-sided empirical test, BH correction across 35 tests per group); open circles, not significant. A thick connector denotes a robust control minus ALS difference.

The strongest differences were in ALS Consortium lumbar spinal cord. Central dogma, OXPHOS and metabolism genes were shifted below random genes in both groups but further in ALS (ALS shifts -0.088 to -0.163; control shifts -0.036 to -0.083; all P_FDR_ < 0.005). This gave robust case-control differences of +0.052, +0.080 and +0.078 (all P_FDR_ < 0.001), which held in 80-96% of draws with cases subsampled to 50 controls (median differences +0.054 to +0.077). Protein import showed a similar difference (+0.055, P_FDR_ = 0.030) but reached P_unadj_ < 0.05 in only 54% of draws.

ALS Consortium cervical spinal cord showed the same pattern with smaller effects. OXPHOS (+0.041, P_BH_ = 0.002) and metabolism (+0.035, P_FDR_ < 0.001) differed robustly, holding in 80% and 86% of subsampled draws. OXPHOS genes were the most strongly shifted of any pathway in this tissue, in both controls (-0.127) and ALS (-0.168). Dynamics C surveillance (+0.051, P_FDR_ = 0.002) was significant but power-limited (72% of draws), and protein import was significant only under the unmatched null.

In ALS Consortium motor cortex, OXPHOS and central dogma genes were shifted below random genes to a similar extent in both groups (OXPHOS: controls −0.113, ALS −0.121; difference +0.007, P_FDR_ = 0.81). Instead, the robust difference was in metabolism (+0.071, P_BH_ < 0.001; 96% of draws). Here, ALS metabolism genes were shifted below random (−0.054, P_FDR_ < 0.001) while control metabolism genes were not (+0.017, P_FDR_ = 0.37). Dynamics C surveillance showed the same pattern (+0.091, P_FDR_ = 0.002), but held in only 70% of draws, consistent with the small control group (n = 32).

In the KCL Brain Bank, differences were smaller. Metabolism again differed robustly (+0.031, P_FDR_ = 0.002; 84% of draws), with ALS metabolism genes shifted below random (−0.028, P_FDR_ = 0.003) and control genes not (+0.003, P_FDR_ = 0.72). OXPHOS (+0.041, P_FDR_ = 0.011) and protein import (+0.052, P_FDR_ = 0.018) were significant under both nulls but held in only 64% and 62% of draws. Signalling (+0.054) was significant only under the unmatched null (P_FDR_ = 0.041 unmatched, 0.074 matched), reflecting a positive shift in controls (+0.061, P_FDR_ = 0.015).

Answer ALS iPSC-derived motor neurons showed the opposite direction to post-mortem tissue. OXPHOS (-0.060, P_FDR_ < 0.001) and dynamics C surveillance (-0.055, P_FDR_ = 0.002) differed robustly, holding in 92% and 86% of draws with cases subsampled to 152. In both pathways, control genes were shifted below random genes (-0.063 and -0.090, P_FDR_ = 0.004 and 0.002) while ALS genes were not (-0.004 and -0.035, P_FDR_ = 0.82 and 0.12). Central dogma (-0.023) and small molecule transport (-0.044) were significant but power limited. Expression matching had the largest effect in this dataset. Null means moved by 0.006-0.040, compared with less than 0.009 in post-mortem tissues. Protein import and metabolism lost significance after matching, but the robust OXPHOS and dynamics C surveillance differences remained.

## Discussion

This study used genetic, transcriptomic and co-expression approaches to ask which mitochondrial pathways are linked to ALS, whether they act upstream or downstream of disease, and in which patients they may play a more important role.

Three main findings emerge. First, inherited ALS risk is modestly enriched in mitochondrial translation, localising mostly to the mitoribosome, with a nominal enrichment in ROS and glutathione metabolism that is largely attributable to *SOD1*. Second, in end-stage tissue, mitochondrial genes are broadly upregulated in ALS motor cortex. This is largely a global shift across the mitochondrial genes, but OXPHOS subunits, and in motor cortex the mitoribosome, are enriched beyond it. OXPHOS subunits ranked above other mitochondrial genes in every dataset, including iPSC-derived motor neurones. Spinal cord shows little change relative to the transcriptome as a whole, but within the mitochondrial genes, central dogma genes are relatively depleted and detoxification genes enriched. At the gene level, genetically predicted expression and differential expression did not agree in direction more often than expected by chance. Third, cell-type adjusted mitochondrial pathway activity separated individuals only weakly into low and high activity groups (mean silhouette 0.22-0.33). In motor cortex these groups were associated with ALS diagnosis in both cohorts, and in lumbar spinal cord the high activity group had longer disease duration (adjusted HR 0.69, P = 0.013). However, no individual pathway was associated with survival after correction in adjusted models. Taken together, the results implicate mitochondrial translation at two levels, through inherited variation that contributes to risk and through expression changes in affected tissue, but this study was not able to provide evidence that the two are directly linked.

Thirty-one mitochondrial central dogma genes reached nominal gene-level significance in MAGMA. They included rRNA and tRNA modification enzymes (*MRM1*, *NSUN3*, *DUS2*, *PUS1*), five mt-tRNA synthetases, the translation factors *TUFM* and *MTIF2*, and proteins from both the small and large mitoribosomal subunits. Rather than a single strong locus, this suggests a signal distributed across a functional pathway, in keeping with the polygenic architecture of ALS (Van Rheenen *et al*., 2021). TWAS and SMR results add gene-level direction to this. Higher predicted expression of *MRPL28*, *MRPL48* and *MRPL38* was associated with increased risk and lower predicted expression of *MRPS33*, *NARS2* and *DUS2* was associated with increased ALS risk. Colocalisation support was reasonable for DUS2 (PP4 up to 0.85) but weak for *MRPS33* (PP4 ≤ 0.34), so the *MRPS33* association may still reflect linkage with a nearby causal variant. A recent Mendelian randomisation analysis reported higher genetically predicted *MRPS33* expression being linked to lower ALS risk (although without evidence of colocalization for this gene), a result consistent with the finding reported here (Du *et al*., 2025). In ALS motor cortex, *MRPS33* expression moved in the opposite direction to that predicted by genetics. It was higher in ALS motor cortex in our cell-type-adjusted models, though not significantly and was significantly higher in an analysis of the same ALS Consortium data that did not adjust for cell composition and included other technical covariates (log2FC = 0.23, P_FDR_ = 0.0007; Humphrey *et al*., 2026). This suggests that part of the increase reflects differences in cell composition, but opposite directions within a single complex may point to the importance of subunit stoichiometry rather than overall expression. The mitoribosome comprises ∼80 proteins (Amunts *et al*., 2015) that are incorporated through a coordinated sequence of early and late binding (Bogenhagen *et al*., 2018), so altered supply of individual subunits could affect assembly and function in different ways. Genetically predicted expression results should, however, be read as annotation rather than replication, because they derive from the same GWAS and the genes were selected on their MAGMA signal. The *MRPL28* association also lacked colocalisation support, so it may reflect linkage with a nearby causal variant.

Enrichment of ALS risk was also nominally detected in ROS and glutathione metabolism (P_unadj_ = 0.001). *SOD1* was the top gene in both this 25 gene set and its parent detoxification pathway, so this result largely reflects recovery of an established ALS gene. In a set of this size, a single strongly associated gene can drive set level significance, and the result does not, on its own, implicate oxidative stress handling more broadly. Repeating the test without *SOD1* would address this. *SOD1* also showed no consistent TWAS or SMR support, in line with *SOD1*-mediated risk acting mainly through protein misfolding and toxic gain of function rather than altered expression (Vande Velde *et al*., 2008; Israelson *et al*., 2010). *SOD1* was however, more highly expressed in ALS motor cortex (ALS Consortium P_FDR_ = 0.004; KCL Brain Bank P_unadj_ = 0.008), but not in spinal cord or iPSC-derived motor neurones. This upregulation may be part of the broader increase in mitochondrial gene expression in ALS cortex.

Genetically predicted expression and differential expression in ALS tissue were largely independent. Across all MitoCarta3.0 genes, the two agreed in direction for 48-52% of genes, no more often than expected by chance, and agreement did not improve when restricted to genes with nominal genetic associations. For example, lower predicted *NARS2* expression was associated with risk, yet *NARS2* was nominally more highly expressed in ALS motor cortex and lumbar spinal cord. One interpretation is that end stage expression changes are driven by the disease process rather than by inherited regulatory variation, as would be expected if they are downstream of disease. However, this conclusion is limited by the set-up of this test. For each gene we used the TWAS or SMR panel with the strongest association, which could be blood or a brain region whose regulatory effects may not apply in motor cortex or spinal cord. Genetically predicted effects on expression are typically small relative to disease-associated changes in tissue, and TWAS associations can be driven by co-regulated neighbouring genes rather than the gene tested. Finally, the binomial and correlation tests treated genes as independent, whereas mitochondrial genes are strongly co-expressed, so the effective number of independent observations is smaller than the number of genes, and a permutation test that preserves this correlation structure would give a more appropriate null. The lack of concordance should therefore be read as an absence of evidence for a link rather than evidence that none exists.

In end-stage tissue, the clearest change was higher expression of mitochondrial genes in ALS motor cortex, seen consistently in two independent cohorts. Most of this was not pathway-specific. When pathways were compared against other mitochondrial genes, significant results fell from 30 to none in the KCL Brain Bank and from 21 to four in ALS Consortium motor cortex. OXPHOS subunits and the mitoribosome remained enriched against this background in ALS Consortium motor cortex, suggesting that these components rise beyond the general mitochondrial shift. Because these models were adjusted for deconvolved cell-type proportions, the shift is unlikely to simply reflect neuronal loss. However, deconvolution estimates are imperfect, and residual confounding by composition or cell state, such as reactive gliosis, cannot be excluded. Higher expression of OXPHOS and mitoribosomal genes could also reflect a compensatory response to energetic stress in surviving cells. Spinal cord, by contrast, showed no enrichment against the genome-wide background, and in cervical cord the central dogma and mtRNA metabolism were depleted, and detoxification enriched, relative to other mitochondrial genes. One possibility is that more extensive loss of motor neurones in end-stage spinal cord leaves less of the affected cell population in which a response could be detected.

iPSC-derived motor neurones lack the neuronal loss and gliosis of end-stage tissue and resemble immature rather than aged neurones (Workman et al., 2023). OXPHOS subunits were the top-ranked pathway in this model and remained enriched against a mitochondrial background, while mitochondrial metabolism as a whole was depleted. Because differences in composition are smaller in this system, the OXPHOS signal could point to an earlier, cell-intrinsic change, although it may also reflect variable differentiation between lines, which immunostaining-based covariates capture only partly.

GSVA scoring separated samples into low and high activity clusters in every tissue, but separation was weak (mean silhouette 0.22-0.33), and in lumbar spinal cord and iPSC-derived motor neurones the two clustering methods agreed poorly. In the KCL Brain Bank, clusters remained associated with RIN and the first surrogate variable even though both had been regressed out. Because diagnosis was protected during regression, any part of these variables that is correlated with diagnosis is retained, so the association between clusters and diagnosis in this cohort may be partly technical. In exploratory survival analyses, neither cluster membership nor individual pathways were consistently associated with survival. The one cluster association was in lumbar spinal cord, where donors in the high activity cluster had longer disease duration (HR 0.62; adjusted HR 0.69, P_unadj_ = 0.013). This result should be interpreted cautiously as the continuous mean activity score was not significant after adjustment for sex and age at onset. Higher organelle contact site activity was nominally associated with longer survival in KCL Brain Bank motor cortex (adjusted HR 0.78 per SD) and ALS Consortium cervical spinal cord (adjusted HR 0.85 per SD), but did not pass correction. This pathway includes *RMDN3* (*PTPIP51*), which partners with *VAPB* to tether the endoplasmic reticulum (ER) to mitochondria, and disruption of ER-mitochondria contacts has been reported in several ALS models (Gomez-Suaga *et al*., 2022). More broadly, post-mortem tissue reflects end stage disease, and its molecular state may say little about the processes that determined the rate of progression. Tissue or cells sampled earlier in disease would provide a better test.

Co-expression between nuclear-encoded mitochondrial genes and mtDNA-encoded genes also differed between pwALS and controls. In all four post-mortem tissues, nuclear-encoded pathway genes were more negatively co-expressed with mtDNA-encoded genes in ALS than in controls, relative to random genes. The difference was strongest in lumbar spinal cord, where central dogma, OXPHOS and metabolism genes were shifted below random genes in both groups but further in ALS, and was weaker in cervical cord, where it was seen for OXPHOS and metabolism. In both motor cortex cohorts, the robust difference was in metabolism, where ALS genes were shifted below random genes and control genes were not. These differences were estimated after adjustment for cell-type composition and were robust both to subsampling of cases and to expression-matched null gene sets.

Negative nuclear-mtDNA co-expression in brain has been described previously and has been suggested to reflect tight regulation of OXPHOS in energetically demanding neurons, matching energy supply to demand while limiting ROS production (Fairbrother-Browne *et al*., 2021). One possibility is a feedback relationship, in which nuclear-encoded machinery is induced when mtDNA-encoded output is low, or vice versa. Under this interpretation, the stronger anti-correlation in ALS post-mortem tissue would indicate tighter, possibly compensatory, coupling between the two genomes in disease, which would fit with the higher expression of nuclear-encoded OXPHOS and mitoribosomal genes in motor cortex. Alternatively, because these are correlations across samples, they may reflect variation in disease stage or residual cell state between individuals with ALS rather than altered regulation within cells. iPSC-derived motor neurones showed the opposite direction, with OXPHOS and dynamics and surveillance genes anti-correlated with mtDNA-encoded genes in controls but not in ALS. If anti-correlation reflects active regulation, this would suggest that coupling is weaker in ALS motor neurones at an immature stage and strengthens, or is selected for in surviving cells, later in disease. These interpretations remain speculative, and expression matching had the largest effect on null distributions in the iPSC dataset.

## Limitations

The main constraint was the use of bulk RNA-seq, which cannot attribute changes to individual cell types. Although differential expression, enrichment, GSVA and co-expression analyses were adjusted for estimated cell-type proportions, these were derived from deconvolution or, in Answer ALS, from immunostaining, and residual confounding by composition or cell state is likely. Post-mortem tissue captures the end of the disease and cannot establish the order of events, and iPSC-derived motor neurones model an immature state and vary in differentiation. GWAS data were limited to European ancestry, the TWAS and SMR results derive from the same GWAS as the MAGMA analysis, and eQTL panels came mainly from blood and bulk brain regions, with limited representation of motor cortex and spinal cord. Control groups were small in some datasets, notably ALS Consortium motor cortex (n = 32), limiting power for case-control comparisons. Finally, survival analyses relied on disease duration in donors who had all died.

## Conclusion

In conclusion, genetic and transcriptomic evidence point to mitochondrial translation in ALS, but at different levels and without a clear link between them. Inherited ALS risk shows a modest enrichment centred on mitochondrial translation and the mitoribosome, with gene-specific directions of effect. In end-stage motor cortex, OXPHOS and mitoribosomal genes are more highly expressed in ALS, and nuclear-mitochondrial co-expression is altered across post-mortem tissues, but these changes are largely unrelated to genetically predicted expression and may represent responses to disease. The OXPHOS signal in iPSC-derived motor neurones warrants follow-up as a possible earlier change. Distinguishing inherited susceptibility from changes that arise during disease will be important when selecting mitochondrial targets for therapy, since targeting a change that is secondary, or that reflects altered cell composition, may be ineffective or even counterproductive.

## Acknowledgments

The authors acknowledge use of the King’s Computational Research, Engineering and Technology Environment (CREATE) (https://create.kcl.ac.uk), which is delivered in partnership with the National Institute for Health and Care Research (NIHR) Biomedical Research Centres at South London and Maudsley and Guy’s and St. Thomas’ NHS Foundation Trusts and part-funded by capital equipment grants from the Maudsley Charity (award 980) and Guy’s and St. Thomas’ Charity (TR130505). We are grateful to the London Neurodegenerative Diseases Brain Bank, and would like to thank the NYGC Consortium for providing the RNA sequencing files. All NYGC ALS Consortium activities are supported by the ALS Association and the Tow Foundation. Data used in the preparation of this article were obtained from the ANSWER ALS Data Portal (AALS-01184). Clinical data and biosamples used in the preparation of this article were obtained from the Answer ALS Foundation Program, ‘Answer ALS’. For up-to-date information on the study, visit https://dataportal.answerals.org.

## Supplementary Figures

**Figure S1.**
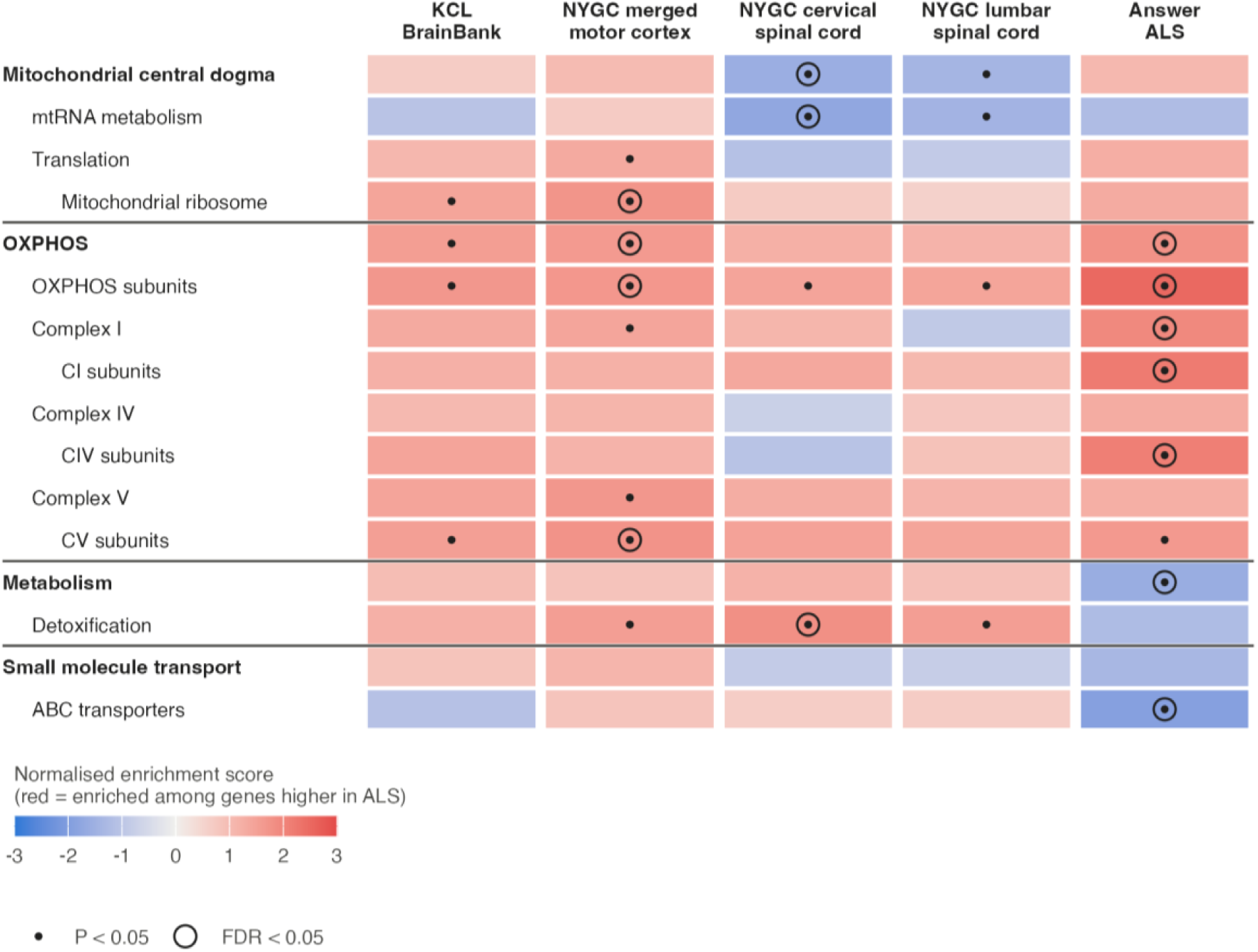
Mitochondrial pathway enrichment in ALS against a mitochondrial gene background.

**Figure S2.**
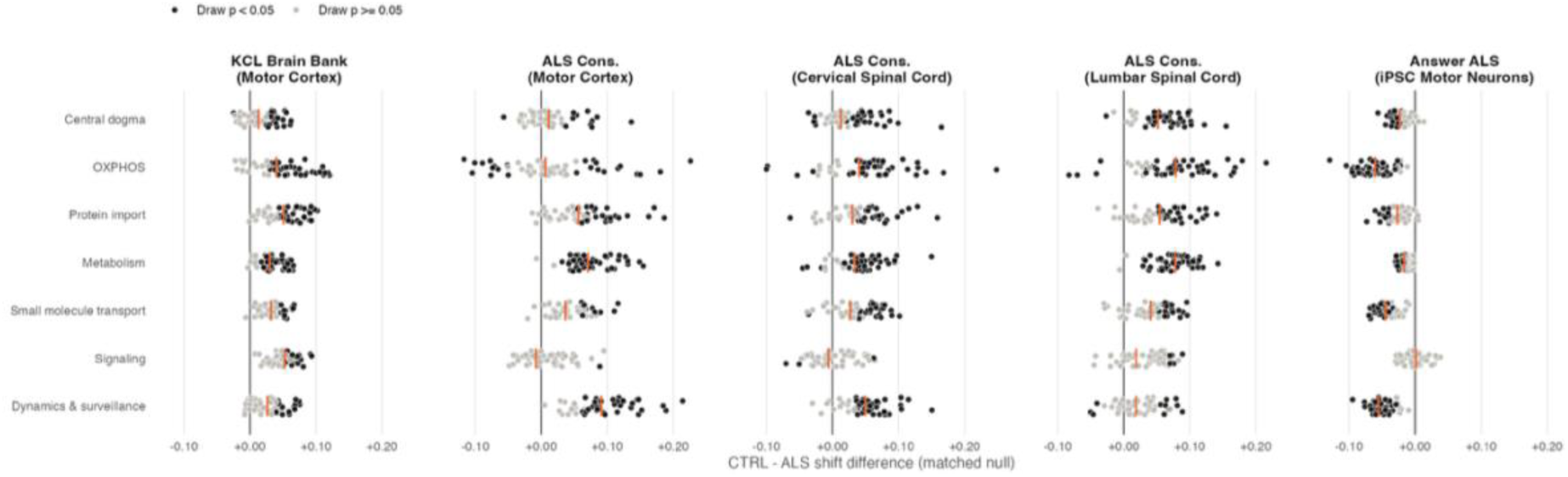
Stability under case subsampling. Each point is the case–control difference from one of 50 draws in which ALS cases were subsampled without replacement to the number of controls (59, 32, 50, 50 and 163). Black points, p < 0.05 in that draw; grey, p ≥ 0.05. Orange ticks mark the full-sample estimate.

